# In-host population dynamics of *M. tuberculosis* during treatment failure

**DOI:** 10.1101/726430

**Authors:** Roger Vargas, Luca Freschi, Maximillian Marin, L. Elaine Epperson, Melissa Smith, Irina Oussenko, David Durbin, Michael Strong, Max Salfinger, Maha Reda Farhat

## Abstract

**Background:** Tuberculosis (TB) is a leading cause of death globally from an infectious agent. Understanding the population dynamics of TB’s causative agent *Mycobacterium tuberculosis* (Mtb) in-host is vital for understanding the efficacy of antibiotic treatment. Here we use longitudinally collected clinical Mtb isolates that underwent Whole-Genome Sequencing (WGS) from the sputa of 307 subjects to investigate Mtb diversity during the course of active TB disease.

**Methods and findings:** We excluded cases suspected of reinfection or contamination to analyze data from 200 subjects, 167 of which met microbiological criteria for delayed culture conversion, treatment failure or relapse. Using technical and biological replicate samples, we defined an allele frequency threshold attributable to in-host evolution. Of the 167 subjects with unsuccessful treatment outcome, 27 (16%) developed new resistance mutations between sampling with 20/27 (74%) occurring in patients with pre-existing antibiotic resistance. Low abundance resistance variants at a purity of ≥19% in the first isolate predicts fixation of these variants in the subsequent sample with 27.0% sensitivity and 95.8% specificity. We identify significant in-host variation in seven genes associated with antibiotic resistance and twenty other genes, including metabolic genes and genes known to modulate host innate immunity by interacting with TLR2. We confirm Rv0095c, Rv1944c, *PPE18, PPE54* and *PPE60* to be under positive selection by assessing phylogenetic convergence across a global and genetically diverse independent sample of 20,352 isolates.

**Conclusions:** Our large sample provides a comprehensive picture of the mutational dynamics *in-host* during active TB disease. We demonstrate a framework to study temporal changes in Mtb population diversity using average depth WGS data. We show that minor variants can be used to inform antibiotic treatment regimens in patients with TB. Furthermore, we detect a signature of positive selection in-host, possibly stemming from innate immune pressure and informing our understanding of host-pathogen interactions.

## INTRODUCTION

Tuberculosis (TB) and its causative pathogen *Mycobacterium tuberculosis* (Mtb) remain a major public health threat (World Health Organization 2018). Yet the majority of individuals exposed to Mtb clear or contain the infection, and only 5-10% of those infected develop active TB disease at some point in their lifetime (Pai et al. 2016). While basic human immune mechanisms to Mtb have been identified, attempts at effective vaccine development guided by these mechanisms have repeatedly failed (Ernst 2018b). Global efforts that include scale up of directly observed therapy have also been challenged by rising estimates of multidrug resistance. Mtb is an obligate human pathogen (Gagneux 2018). Infection and disease involves a complex human host-pathogen interaction that is both physically and temporally heterogeneous (Lin et al. 2014). Consequently all selective forces acting on Mtb will originate within the host, and the study of temporal dynamics of this is likely to inform antibiotic treatment (Sun et al. 2012) and rational vaccine design (Ernst 2018b).

At long timescales, signatures of positive selection associated with antibiotic resistance have been characterized, but epitope regions appear to be under purifying selection (Farhat et al. 2013; Coscolla et al. 2015; Brites and Gagneux 2015; Comas et al. 2010) calling into question how Mtb interacts with host adaptive immunity. Little is known about selection at short timescales, such as within single infections. Drug pressure may select for resistance-conferring mutations, thus an understanding of how the frequency of minor alleles changes longitudinally can inform optimal drug treatment (Sun et al. 2012; Zhang et al. 2016; Didelot et al. 2016). A recent study found treatment relapse to be strongly associated with bacterial factors (Colangeli et al. 2018); therefore there is a need to better characterize these as predictors of treatment response. Bacterial factors of interest include not only low frequency resistance variants but also variants that may induce other phenotypes, such as drug tolerance or more effective immune evasion (Ernst 2018a). To elucidate these processes, we aimed to study how genomic diversity arises in-host in Mtb populations, employing a longitudinal sampling scheme from patients with active TB disease.

The application of genome sequencing technologies to Mtb isolates cultured from clinical samples has highlighted that infection consists of populations of Mtb bacteria rather than single clones devoid of diversity (Marvig et al. 2015; Lieberman et al. 2011, 2014; Copin et al. 2016; Didelot et al. 2016). Differences in observed allele frequencies captured using genome sequencing (**Fig. 1A**) may represent a difference in the genetic composition of the infecting population, commonly referred to as heterogeneity. Mtb population heterogeneity might be present within a host because (1) the host is infected with multiple strains or is re-infected by a new strain (consistent with mixed infection or re-infection) or (2) genetic diversity arises within the Mtb population during infection due to selection or drift (Ford et al. 2012; Lieberman et al. 2016; Guerra-Assunção et al. 2014). However, non-uniform sampling (Trauner et al. 2017), selection during the *in vitro* culture process (Trauner et al. 2017), laboratory contamination (Goig et al. 2020; Wyllie et al. 2018), sequencing error and mapping error all represent examples of experimental error that give rise to heterogeneity of low significance to host-pathogen interactions.

**Fig. 1.**
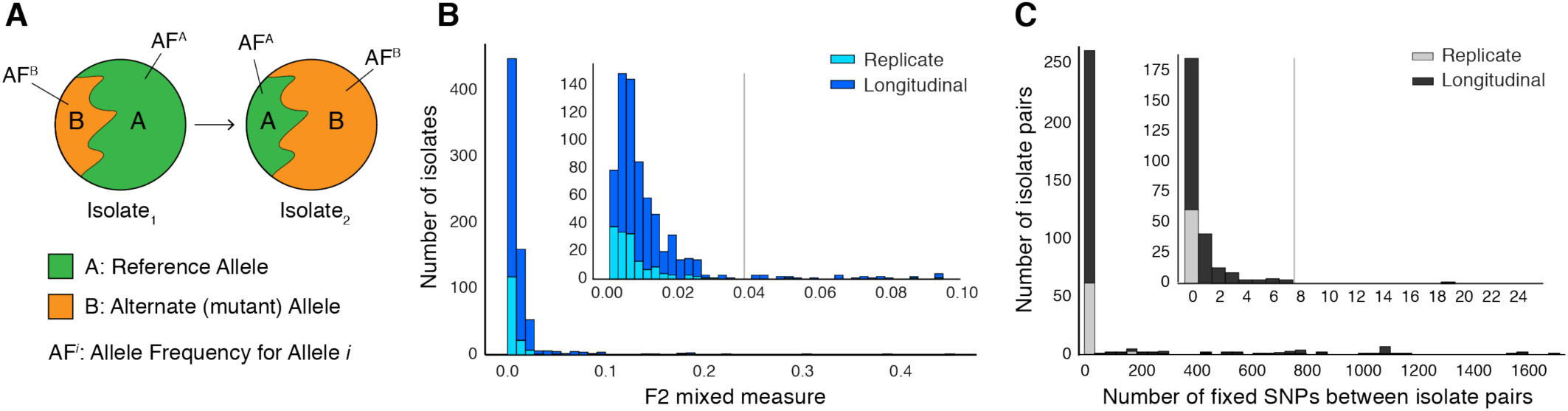
Selection of patients with longitudinal clonal infection. (**A**) Allele frequency change between paired isolates 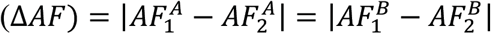. (**B**) The F2 measure >0.04 (**Methods**) was used to identify and exclude isolate pairs with evidence for mixed strain growth at any time point. (**C**) Replicate and longitudinal pairs with fixed SNP (fSNP) distance of >7 were excluded. For longitudinal isolates fSNP>7 was assessed as consistent with Mtb reinfection with a different strain.

Here, we present a framework to overcome these barriers and demonstrate the use of longitudinally collected isolates, pooled sweeps of colonies cultured from sputa, to investigate true in-host diversity with implications for Mtb treatment. We analyzed 614 paired longitudinal isolates representing 307 subjects from eight studies (Casali et al. 2016; Walker et al. 2013; Trauner et al. 2017; Guerra-Assunção et al. 2014; Bryant et al. 2013; Witney et al. 2017; Xu et al. 2018). Many subjects, despite undergoing treatment, remained culture positive at two months intervals or longer meeting microbiological criteria for delayed culture conversion, treatment failure or relapse (**Table S1**). Our sample consisted of 167 subjects fulfilling these criteria, which allowed us to overcome the small sample size problem present in prior studies. We find a high turnover of low-frequency alleles in loci associated with antibiotic resistance but that mutant alleles in these loci that rise to a frequency of 19% are predicted to fix in-host with a sensitivity of 27.0% and specificity of 95.8%, a proof of concept that WGS can aide in predicting resistance amplification. We show that changes in allele frequency are common among replicate isolates and that changes in frequency of 70% are indicative of in-host evolution using archived MTB isolates. We demonstrate that many loci involved the acquisition of antibiotic resistance and modulation of innate host-immunity appear to be under positive selection.

## RESULTS

### Identifying clonal Mtb populations in-host

To isolate the *in vivo* clonal dynamics of Mtb during infection among the 307 subjects with longitudinal samples collected during or after antibiotic treatment (**Table S1-S2**), we excluded 32 subjects with isolate microbiological contamination at any time point (Goig et al. 2020), and 31 subjects with evidence for mixed infection with two or more Mtb lineages (Wyllie et al. 2018) (**Fig. 1B** and **Fig. S1**). We also excluded 44 subjects with evidence for re-infection with a different Mtb strain between the first and second time points, using a pairwise genetic distance >7 fixed SNPs (fSNPs) (**Methods, Fig. 1C** and **Fig. S1**). We implemented WGS SNP calling filters to minimize the likelihood of false positives and estimated the error rate of our analysis pipeline using a control dataset of 82 isolate pairs (162 total) that were *in vitro* technical or biological replicates (**Methods, Fig. 4** and **Fig. S1**). Of the 307 subjects, 200 had isolate pairs that passed all filters (representative of clonal infection) (**Table S3**), with an estimated false positive SNP rate of 0.0527 or less. The 400 isolates represented the five main Mtb lineages.

**Fig. 2.**
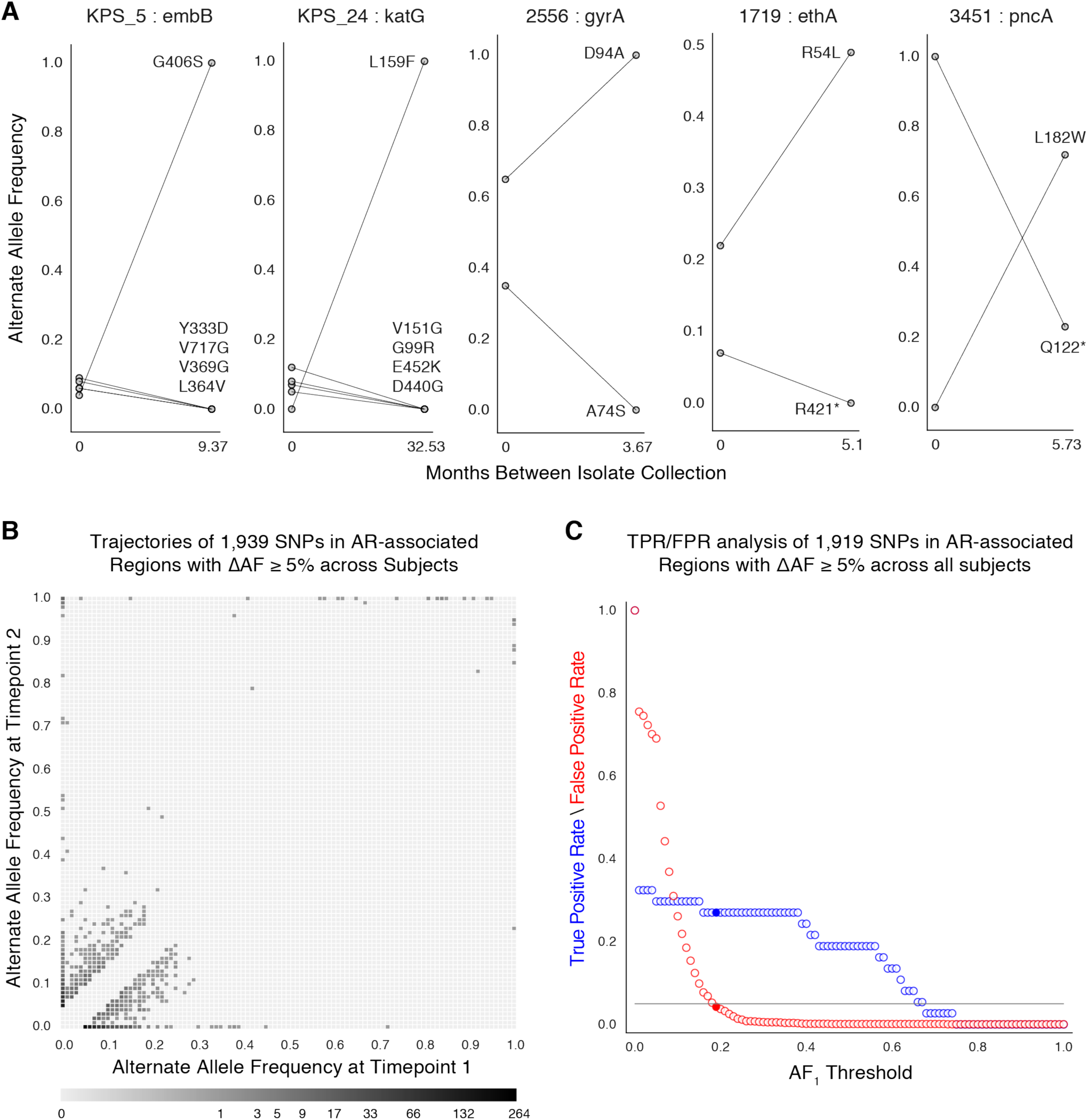
Allele frequency dynamics within antibiotic resistance loci. (**A**) The antibiotic resistance genes *embB, katG, gyrA, ethA*, and *pncA* demonstrate evidence for competing clones during infection (other examples found are displayed in **Fig. S2**). Each mutant allele is labeled with amino acid encoded by the reference allele, H37Rv codon position, and amino acid encoded by the mutant allele. (**B**) The allele frequency trajectories for SNPs that occur in subjects over the course of infection can be used to study the prediction of further antibiotic resistance using the frequency of alternate alleles detected in the longitudinal isolates collected from subjects. (**C**) Plot of true positive rate (TPR) and false positive rate (FPR) for detecting eventual fixation of a resistance allele (*AF*_2_ ≥ 75%) as a function of initial allele frequency (*AF*_1_).

**Fig. 3.**
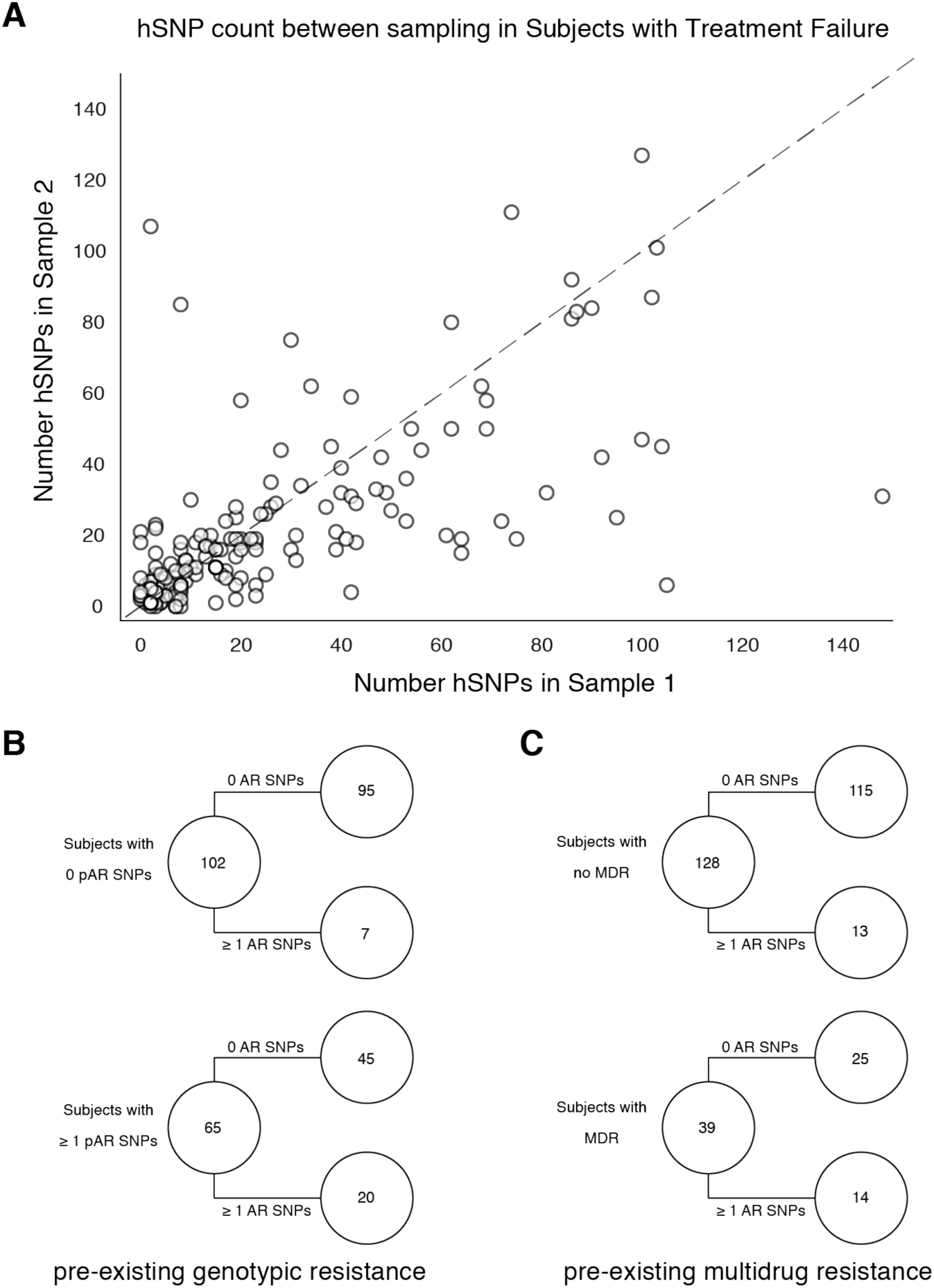
Pre-existing resistance is associated with resistance amplification. We called heterozygous SNPs (hSNP) in each isolate from a subject with clonal infection classified as failing treatment (N=167). We defined hSNPs as a SNP called in an isolate with an alternate allele frequency between 25% and 75% (**Methods**). (**A**) The number of hSNPs called in the 2^nd^ sample isolated vs the number of hSNPs called in the 1^st^ sample isolated from each of 167 subjects. The dashed line is y = x. (**B-C**) Among subjects who fail treatment, (**B**) subjects with pre-existing mutations that confer antibiotic resistance and (**C**) those that have pre-existing MDR are more likely to acquire antibiotic resistance mutations throughout the course of infection.

**Fig. 4.**
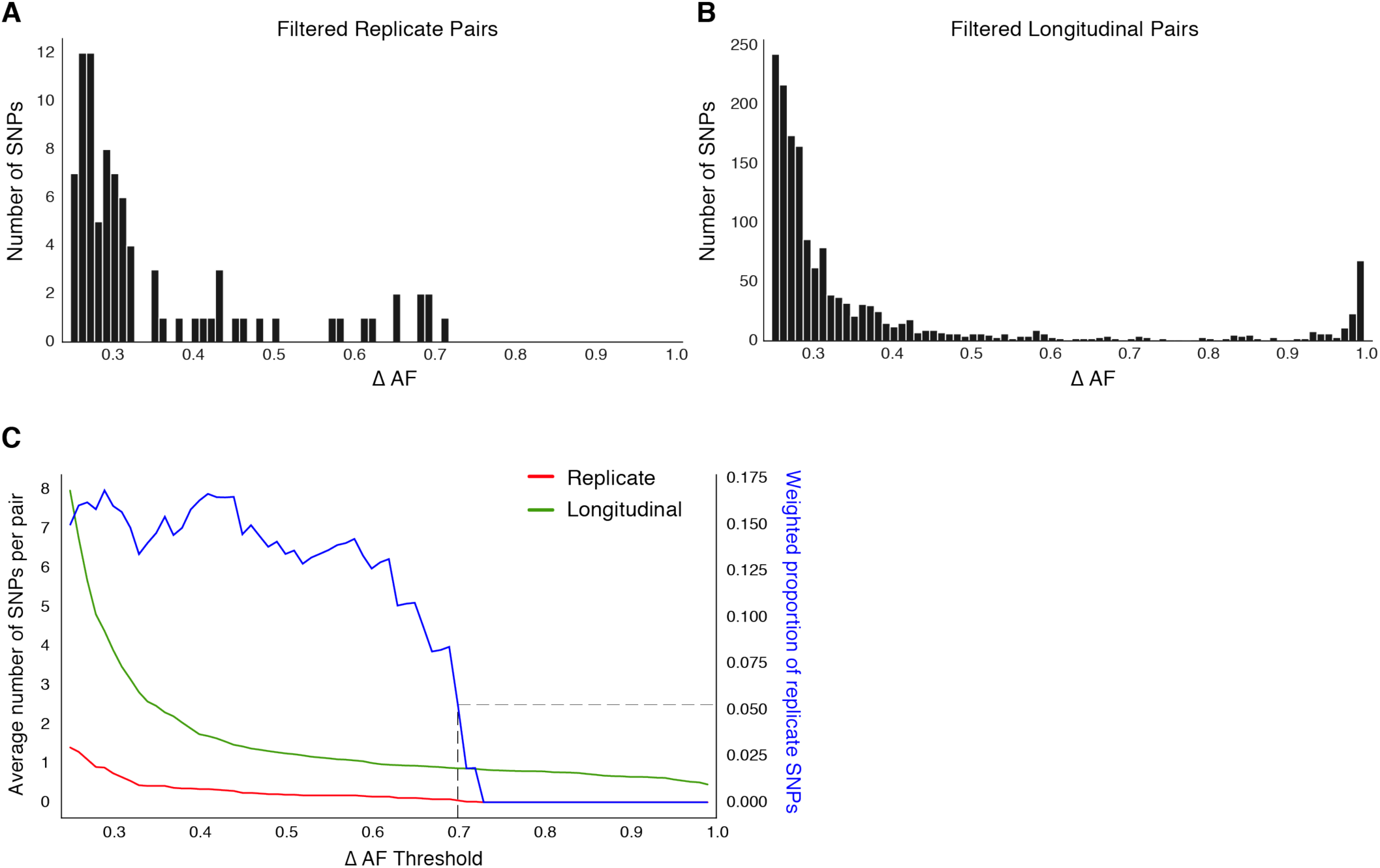
Replicate pairs reveal levels of biological noise associated with repeated sampling. (**A, B**) We analyzed the distribution of ΔAF for all SNPs detected across all replicate pairs (*m* = 62) and longitudinal pairs (*n* = 200) for SNPs where ΔAF ≥ 25%. (**B**) SNPs were detectable at lower levels of ΔAF for both types of isolate pairs, but SNPs with higher values of ΔAF were only found in longitudinal pairs. (**C**) To determine a ΔAF threshold for calling SNPs representative of changes in bacterial population composition in-host, we calculated the average number of SNPs per pair of isolates at different ΔAF thresholds for both replicate and longitudinal pairs. At a ΔAF threshold of 70% the number of SNPs between replicate pairs represents 5.27% of the SNPs detected amongst all replicate and longitudinal pairs, weighted by the number of pairs in each group.

### In-host pathogen dynamics in antibiotic resistance loci

Resistance mutations found at low frequencies in-host may indicate the impending development of clinical resistance (Sun et al. 2012; Trauner et al. 2017; Zhang et al. 2016), To investigate temporal dynamics related to antibiotic pressure, we identified non-synonymous and intergenic SNPs within a set of 36 predetermined resistance loci associated with antibiotic resistance (Farhat et al. 2013, 2016) (**Table S5**) that changed in allele frequency by at least 5% (Sun et al. 2012) between the first and second sampling time point (**Methods**). We detected 1,939 such SNPs across our sample of 200 subjects (**Fig. 2B**), 1,774 were non-synonymous, 91 were intergenic, and 74 occurred within the *rrs* region (**Table S6**).

We searched for signs of selection by identifying clonal interference, or evidence of competition between strains with different drug resistance mutations (Trauner et al. 2017; Sun et al. 2012; Zhang et al. 2016). We characterized this in longitudinal isolates fulfilling three criteria: (i) isolates containing multiple resistance SNPs in the same gene within the same subject, (ii) at alternate allele frequencies that change in opposing directions over time and (iii) the alternate (mutant) allele frequency was intermediate to high at ≥ 40% in at least 1 isolate (Farhat et al. 2016) for at least one of the co-occurring SNPs. This identified 11 cases of clonal interference (**Fig. 2A** and **Fig. S2**), demonstrating most often the fixation of a single allele in the second isolate from a mixture of multiple alleles at lower frequencies in the first isolate collected.

### Allele frequency >19% predicts subsequent fixation of resistance variants

We determined the lowest AR allele frequency that can accurately predict the development of fixed resistance alleles later in time (Sun et al. 2012; Zhang et al. 2016). We discarded 20 SNPs that were fixed (*AF* >75%) in both isolates to study the AF trajectories of 1,919 AR SNPs detected with a change in allele frequency, ΔAF ≥ 5% between time points. We calculated the true positive rate (TPR) and false positive rates (FPR) for varying values of *AF*_1_ ∈ {0,1, 2, …, 99,100}% (**Fig. 2C, Methods**). For the purposes of prediction, we allowed a maximum FPR of 5% and found the optimal classification threshold to be 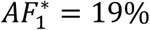 with an associated sensitivity of 27.0% and a specificity of 95.8%. Ten mutant alleles across 14 isolates from 7 subjects had a frequency between 19% and 75% at the first time point and rose to fixation at the second time point (mean ΔAF 41%).

### Determinants of antibiotic resistance acquisition and microbiological treatment failure

All study subjects had either recently completed treatment or were receiving treatment when samples were collected (Walker et al. 2013; Casali et al. 2016; Trauner et al. 2017; Xu et al. 2018; Guerra-Assunção et al. 2014; Bryant et al. 2013; Witney et al. 2017), but details of the treatment regimen were not available (**Table S1**). We identified overall rates of resistance acquisition by focusing on AR SNPs with moderate to high ΔAF ≥ 40% given prior evidence of association between such SNPs and phenotypic resistance (Farhat et al. 2016). Thirty-eight AR SNPs were detected in 195/200 subjects with clonal infection and known isolate sampling date. AR acquisition was more likely as the time between sampling, and likely antibiotic exposure, increased, with the OR of AR acquisition being 1.023 per 30day increment (95% *CI* 1.002, 1.045, *P* = 0.035 Logistic Regression).

We focused on subjects with samples collected ≥2 months apart, by definition culture positive at sample collection time, because they meet microbiological criteria for *delayed culture conversion, failure* or *relapse* (hitherto failure for brevity) (WHO). Of the 269 subjects with mixed or clonal infection and reinfection as well as complete isolate sampling dates, 230 had samples collected ≥2 months apart and consisted of 35 reinfections (15%), 28 mixed infections at one or two time points (12%) and the majority, 167 (73%), had clonal infection (**Fig. S1**).

We examined the relationship between pre-existing resistance and new AR acquisition in clonal treatment failure. Pre-existing resistance was defined as ≥1 fixed AR SNPs in the first isolate (Farhat et al. 2016) (**Methods**). Two hundred thirty pre-existing AR SNPs were identified with 39% (65/167) of failure subjects harboring resistance to any drug at the first sampling (**Fig. 3B, Table S7**). The majority of this resistance was MDR (multidrug resistance to at least isoniazid and a rifamycin), 60% (39/65) (**Fig. 3C**). New resistance acquisition occurred mostly in subjects with pre-existing resistance 20/27 (74%) (*OR* = 6.03, *P* = 6.8 × 10^−5^ Fisher’s exact test) or pre-existing MDR (*OR* = 4.95, *P* = 3.8 × 10^−4^ Fisher’s exact test).

We also quantified genome-wide Mtb diversity in-host among the patients with microbiological treatment failure. We reasoned that as these patients are not on or not adherent to effective antibiotic treatment, their effective pathogen population size may be large and prone to more genetic drift or turnover of minority variants with and without selection (Trauner et al. 2017). We counted the number SNPs with an alternate allele frequency between 25% and 75% (**Methods**) at each time point as a conservative estimate of the number of segregating sites in each population. We found this count to strongly correlate between the first and second time point (**Fig. 3A**) suggesting that minor allele diversity is maintained in-host in patients without effective therapy (median T1=15 hSNPs, median T2=16 hSNPs, *r*^2^ = 0.43, *P* = 9.5 × 10^=−22^ Linear Regression).

### Genome-wide in-host diversity

Beyond antibiotic pressure, selective forces acting on the infecting Mtb population in-host are largely unknown. To investigate this reliably across the entire Mtb genome, we first examined the genome-wide allele frequency distribution for both technical replicates (*in vitro* technical or biological replicates, sample size m=62 after exclusions) and in-host longitudinal pairs (**Fig. 4** and **Fig. S1**). We detected five SNPs in *glpK* (with ΔAF ≥ 25%) among five replicate pairs (mean ΔAF=45%) consistent with an adaptive role for *glpK* mutations *in vitro* (Pethe et al. 2010; Vargas and Farhat 2020) and accordingly excluded this gene from further analysis (**Methods**). The genome-wide AF distribution in both replicate and longitudinal pairs demonstrated an abundance of SNPs with low ΔAF likely resulting from noise or technical factors. To clearly distinguish signal related to in-host factors from noise, we determined the ΔAF threshold above which SNPs/isolate-pair were rare among technical replicates *i.e.* constituted 5% or less of total SNPs (**Fig. 4**). We determined this ΔAF threshold to be 70% and selected 174 SNPs that developed in-host (in-host SNPs) among the 200 TB cases (**Fig. 4C, Table S8**).

### Characteristics of mutations in-host

Of the 174 SNPs, 112 were non-synonymous, 42 synonymous, and 21 were intergenic (**Fig. 5C**). The 153/174 coding SNPs were distributed across 127/3,886 genes and were observed in 71/200 subjects (**Fig. 5B** and **Fig. 5D**). We analyzed the spectrum of mutations and found the GC > AT nucleotide transition to be the most common. The GC > AT transition is putatively due to oxidative damage including the deamination of cytosine/5-methyl-cytosine or the formation of 8-oxoguanine (Ford et al. 2011; Dillon et al. 2015). The transversion AT > TA was the least common substitution (**Fig. 6A**). We expected the number of SNPs detected between longitudinal isolates to increase with time between isolate collection. Regressing the number of SNPs per subject on the timing between isolate collection (for 195 subjects with isolate collection dates) (**Fig. 6B**), we found SNPs to accumulate at an average rate of 0.56 SNPs per genome per year (*P* = 7 × 10^−12^) consistent with prior *in vivo* estimates (Walker et al. 2013; Ford et al. 2011).

**Fig. 5.**
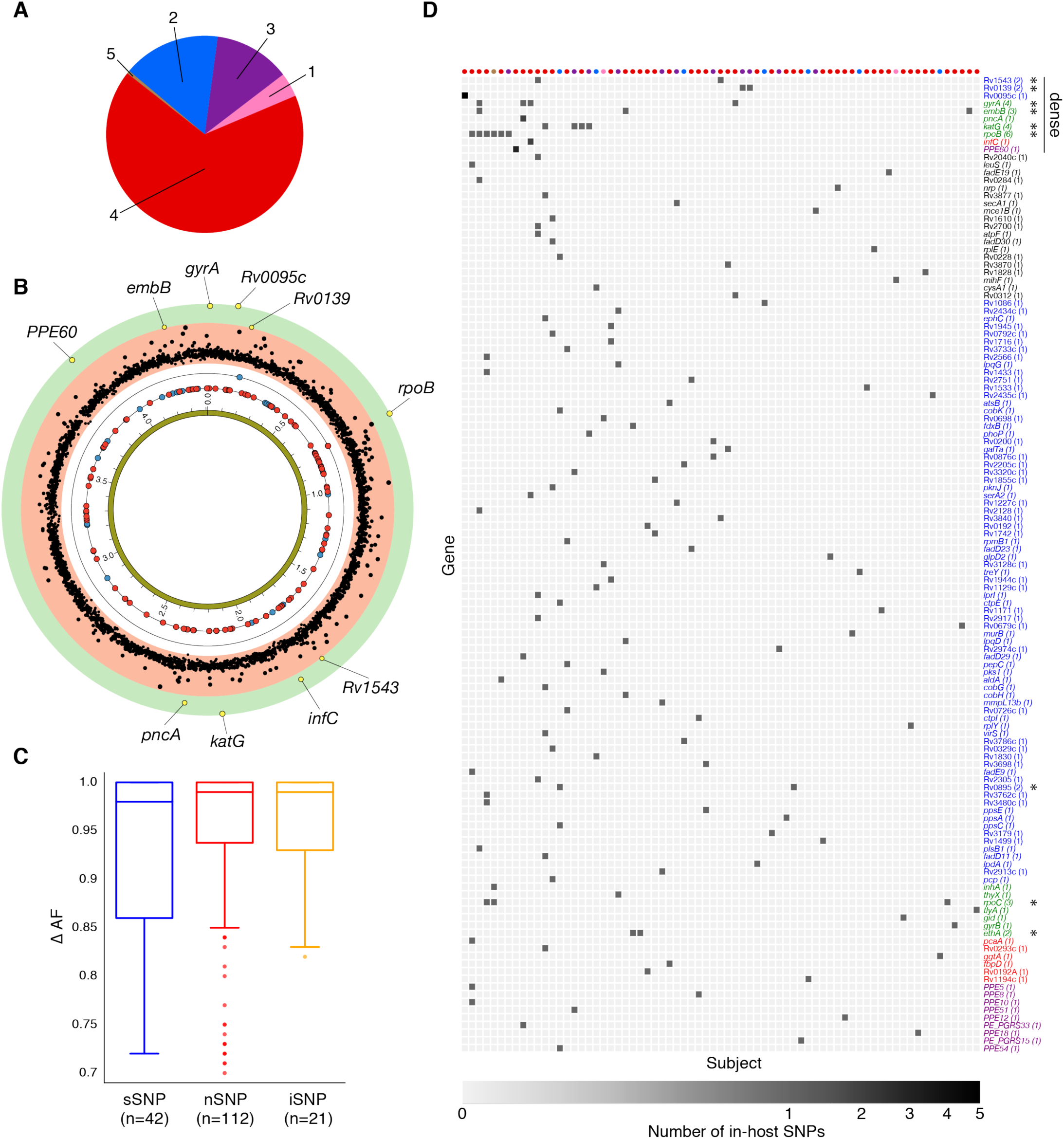
Genome-wide diversity in 200 clonal Mtb infections. (**A**) Distribution of five major Mtb lineages among the 200 clonal Mtb infections. (**B**) Distribution of 153 in-host SNPs within coding regions among the 200 longitudinal isolate pairs across the 4.41 Mbp Mtb genome (blue circles: synonymous, red circles: non-synonymous). Blue and red circles on the innermost black ring indicate the locations of SNPs detected in one patient; circles on the next ring represent SNPs detected in two patients. The −log_10_(p-value) of the mutational density test (**Methods**) by gene is plotted in the outermost, red and green, regions. Labeled yellow circles represent genes significant at the bonferroni-corrected cutoff (*α* = 0.05/3,886). (**C**) Distribution of ΔAF by SNP type: sSNP: synonymous, nSNP: non-synonymous, iSNP: intergenic. (**D**) Heat-map of SNPs per gene (rows) and patient (columns). Colored circles across columns indicate the strain phylogenetic lineage (as represented in **A**). Gene names colored according to gene category (**Fig. 6D**) with parentheses indicating the number of subjects with a SNP in a given gene. *Indicates genes in which SNPs are detected within multiple subjects.

**Fig. 6.**
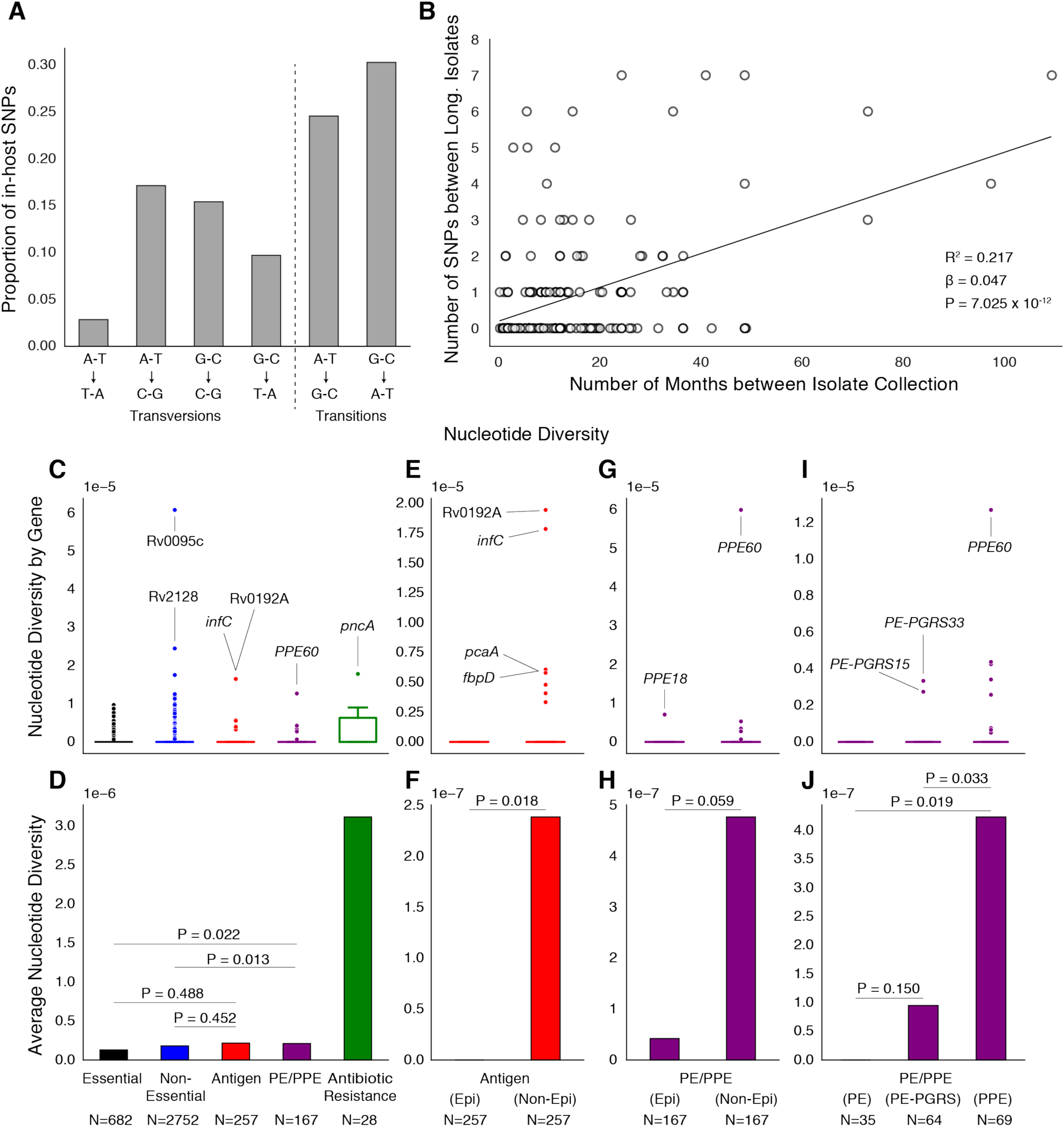
PE/PPE genes vary considerably within host while putative antigens remain conserved. (**A**) Mutational spectrum of in-host SNPs. (**B**) In-host SNP counts vs. time between isolate collection (195/200 subjects with dates shown, *W (Walker et al. 2013) isolates only had year of collection). (**C**) boxplots of nucleotide diversity by gene within each of 5 non redundant categories (see text). (*n* = number of genes). (**D**) Average nucleotide diversity across genes by category. Nucleotide diversity in epitope and non-epitope region (**Methods**) of each gene in the Antigen (**E, F**) and PE/PPE (**G, H**) gene categories. (**l, J**) PE/PPE genes separated into three non-redundant categories: PE, PE-PGRS, and PPE. (**J**) The average nucleotide diversity by category. (**I**) box plot of nucleotide diversity by gene.

### Simulations and PacBio sequencing demonstrate a low false-positive rate in repetitive regions

Several SNPs detected were in the GC-rich repetitive PE/PPE gene family (Brennan and Delogu 2002). Variants called on these genes are commonly excluded from comparative genomic analyses (Coscolla et al. 2015; Comas et al. 2010; Copin et al. 2016; Casali et al. 2016) due to the limitations of short-read sequencing data and the possibility of making spurious variant calls, however the rates at which these false calls occur has not been evaluated. We reasoned that our stringent filtering criteria, quality of sequencing data and depth of coverage allowed us to reliably detect variants in these regions of the genome, with the potential to uncover variation in these understudied regions of the genome.

We took several approaches to test the rate of false-positives for the single base-pair mutations observed in our analysis (**Methods**). First, we introduced the mutant alleles observed in-host (**Table S10**) into a set of Mtb reference genomes belonging to different lineages and simulated short read sequencing data from these modified genomes (**Fig. S5**). We then used our variant calling pipeline to call bases from this simulated data. We observed a high recall rate of the introduced mutant alleles and a very low number of false positive base calls (zero in most cases) within the loci containing modified alleles (**Fig. S6**). Second, we assessed the congruence in variant calls between short-read Illumina data and long-read PacBio data for a set of isolates that underwent sequencing with both technologies (**Methods)**. Unlike Illumina generated reads, PacBio reads are much longer and have randomly distributed error profiles (Rhoads and Au 2015). With high coverage, PacBio sequencing can reliably reconstruct full microbial genomes and identify SNPs in repetitive regions. The comparison with PacBio assemblies confirmed empirically a low rate of false positive base calls in genomic regions where we observed in-host SNPs (**Methods**). Third, we confirmed the five phylogenetically convergent in-host SNPs in PPE genes *PPE18, PPE54* and *PPE60* (see below) through manual inspection of the read alignment (**Fig. S7-S9**).

### Antibiotic resistance and PE/PPE genes vary while antigens remain conserved

To understand how different classes of proteins evolve in-host, we separated Mtb genes into five non-redundant categories (**Methods**). The vast majority of genes in each category did not vary within subjects (**Fig. 6C**). Antibiotic resistance genes were on average the most diverse category while Essential genes varied the least (**Fig. 6D**). Antigen genes appeared to be as conserved as were both Essential (*P* = 0.49 Mann-Whitney U-test) and Non-Essential genes (*P* = 0.45 Mann-Whitney U-test) while PE/PPE genes showed higher levels of nucleotide diversity than both Essential (*P* = 0.022 Mann-Whitney U-test) and Non-Essential genes (*P* = 0.013 Mann-Whitney U-test) (**Fig. 6D**).

### PE/PPE variation is independent of T-cell recognition

To test whether variation in Antigen or PE/PPE genes occurred in response to T-cell recognition, we separated each gene in these categories into (CD4^+^ and CD8^+^ T-cell) epitope and non-epitope concatenates and recalculated nucleotide diversity for these concatenates (**Fig. 6E-H**). For both Antigen and PE/PPE genes (**Fig. 6F** and **Fig. 6H**), epitope concatenates were less diverse than non-epitope concatenates (*P* = 0.018 and *P* = 0.059 respectively, Mann-Whitney U-test). Only one in-host SNP was detected within an epitope-encoding region in the gene *PPE18* (**Fig. 6G** and **Fig. S4, Table S12**). This suggests that T-cell recognition does not drive diversity in these regions. Looking within the three PE/PPE subfamilies (**Fig. 6I-J**) (Brennan 2017), the PPE genes appeared more diverse in-host than PE genes and PE-PGRS genes (*P* = 0.019 and *P* = 0.033 respectively, Mann-Whitney U-test).

### Identifying candidate pathoadaptive loci from genome-wide variation

To identify genes involved in pathogen adaptation (Lieberman et al. 2011; Marvig et al. 2015), we applied a test of mutational density (Farhat et al. 2014) (**Methods**) by pooling variation across all 200 pairs of genomes and identifying those genes with more mutations than expected under a neutral model of evolution where variants are Poisson distributed across the genome (Farhat et al. 2014) (**Fig. 5B, Table S13, Methods**). We also searched for evidence of convergent evolution *i.e.* genes or pathways where in-host SNPs developed in ≥ 2 subjects (**Table S14, S17**). Seven known antibiotic resistance genes (Didelot et al. 2016; Farhat et al. 2013) had significant mutational density (*α* = 0.05, Bonferroni correction) or were convergent across patients: *rpoB, gyrA, katG, rpoC, embB, ethA* and *pncA* (mutated in six, four, four, three, three, two and one subject respectively) (**Fig. 5B** and **Fig. 5D**). Single in-host SNPs occurred in eight additional known resistance loci including three intergenic regions, and in *prpR*, a gene recently implicated with drug tolerance (Hicks et al. 2018) (**Table S8**).

Three genes with unknown function: Rv0139, Rv0895, and Rv1543 were convergent in two subjects each, two of which (Rv0139, Rv1543) had significant mutational density (P<2×10^− 5^) and; three additional genes including *PPE60* displayed significant mutational density (P<2×10^− 5^) (**Fig. 5B, Table S13**). We found evidence for convergence in six pathways not known to result in antibiotic resistance. These pathways are involved with biotin biosynthesis (*fadD23, fadD29*, and *fadD30*), ribosomal large subunit proteins (*rpmB1, rplE*, and *rplY*), glycerolipid and glycerophospholipid metabolism (*aldA* and Rv2974c), ESAT-6 protein secretion (*eccCa1* and *eccD1*), coenzyme B12/cobalamin synthesis (*cobH* and *cobK*) and the uncharacterized pathway CBSS-164757.7.peg.5020 (*fdxB* and *PPE18*) (**Table S17**).

### In-host mutations display phylogenetic convergence across multiple global lineages

We reasoned that pathoadaptive mutations observed to sweep to fixation in-host and not compromise pathogen transmissibility are likely to arise independently within other subjects and in separate geographic regions in a convergent manner (Farhat et al. 2013). We screened a geographically diverse set of 20,352 sequenced clinical isolates belonging to global lineages 1-6 for mutations observed within host in which the alternate (mutant) allele swept over the course of sampling (141/174 in-host SNPs, **Methods, Table S8, S18**). Conservatively, a mutation was characterized as phylogenetically convergent if it was present in isolates from three or more global lineages but not fixed in any lineage (**Methods**). We identified 26/141 in-host SNPs as phylogenetically convergent in our global sample of isolates (**Table S19**). **Fig. 7** and **Fig. S10-11** display the distribution of convergent alleles across the 20,353 isolates using t-Distributed Stochastic Neighbor Embedding (t-SNE) of the pairwise genetic distance matrix (**Methods**). The convergent alleles included the PPE genes *PPE18* (1 site), *PPE54* (1 site) and *PPE60* (3 sites), as well as Rv0095c (2 sites) and Rv1944c (1 site) both conserved proteins of unknown function. In addition to several SNPs in loci associated with antibiotic resistance, *gyrB* (1 site), *gyrA* (2 sites), *rpoB* (4 sites), *rpoC* (3 sites), *inhA* (1 site), *embB* (3 sites) and *gid* (1 site).

**Fig. 7.**
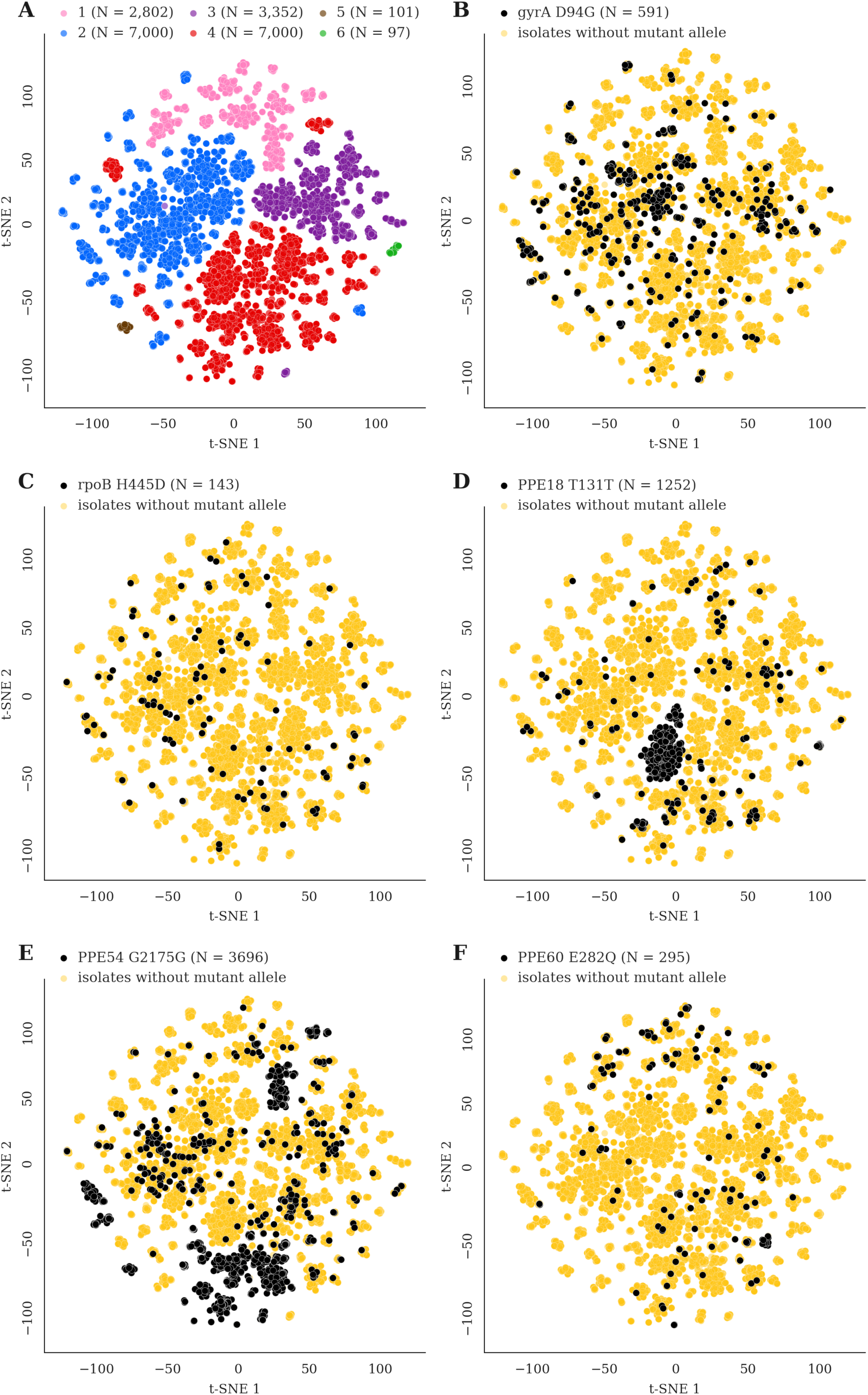
Mutations acquired in-host are phylogenetically convergent. We constructed t-SNE plots from a pairwise SNP distance matrix for our global sample of 20,352 clinical isolates and 128,898 SNP sites (**Methods**). (**A**) Labeling isolates by global lineage revealed that isolates cluster according to genetic similarity. Next, we labeled isolates by whether they carried a mutant allele that was also detected in-host. (**B-F**) Mutations in *gyrA, rpoB, PPE18, PPE54* and *PPE60* were detected in-host (**Table S8**), occur in a global collection of isolates (**Table S18**) and are scattered across the tSNE plots, indicating that they belong to genetically different clusters of isolates (**Table S19**). Furthermore, all mutations with a signal of phylogenetic convergence were detected in isolates belonging to different clusters, confirming that theses mutations must have arisen independently in different genetic backgrounds (**Fig S10-S11**). Each plot is labeled with the gene name each mutation occurs within, amino acid encoded by the reference allele, H37Rv codon position, and amino acid encoded by the mutant allele. N = number of isolates with mutant allele.

## DISCUSSION

In our 400 Mtb populations sequenced from 200 active TB patients heavily enriched for delayed culture conversion, treatment failure and relapse, we find a wealth of dynamics in genetic loci associated with antibiotic resistance, including a high turnover of minor variants (Trauner et al. 2017). Factors that determine negative treatment outcomes are complex and include severity of lung disease, cavitation and adherence to treatment among others (Imperial et al. 2018). Here, we observe that a relatively high percentage of such patients, 16%, develop antibiotic resistance over time. Our findings of a higher rate of resistance acquisition in patients with MDR at the outset and with time between sampling, emphasize the importance of appropriately tailoring treatment regimens as well as close surveillance for microbiological clearance and resistance acquisition by phenotypic or genotypic means. The observed high rate of resistance acquisition also emphasizes Mtb’s biological adaptability and the long duration of drug pressure *in vivo.* Resistance acquisition in the course of one infection is comparatively rare in most pathogenic bacteria (Llewelyn et al. 2017). In addition to clonal acquisition of resistance, we found 27% of patients with unsuccessful treatment outcomes to have mixed infection or reinfection with different Mtb strains. This high percentage suggests that care of these patients and control of disease transmission can be better guided if pathogen sequencing is routinely performed for cases meeting these microbiological criteria especially in high TB prevalence settings where reinfection is more likely.

We provide a proof of concept analysis that minor AR alleles, occurring at a frequency down to 19%, can accurately predict fixation of the variant in >95% of mutations in-host, although we find the sensitivity of this threshold to be low. In the future, higher depth sequencing at fixed sampling durations can elucidate more clearly the role of minor AR allele detection in clinical management of TB treatment.

Various sources of noise contribute to allele frequency changes over time and challenge inference on bacterial composition *in vivo*. Here, we determined an appropriate threshold for identifying mutations in-host using average depth Mtb WGS from cultured isolates and demonstrate the importance of including technical replicate WGS. While culturing sputa *in vitro* enriches Mtb DNA for WGS it also creates experimental noise (Vargas and Farhat 2020). The refinement of methods for DNA extraction directly from sputum (Votintseva et al. 2017), may allow the calling of relevant changes in allele frequencies at lower thresholds in future work. This would permit the unbiased study of loci that may be under frequency-dependent selection, where changes in allele frequencies would unlikely change by as much as 70% as we used here.

We detected 174 alleles rising to near fixation in-host across our sample of 200 subjects. The observed distribution of variants including the high rate of non-synonymous substitutions and the predominance of GC > AT variants are consistent with the hypotheses of purifying pressure on synonymous variants and oxidative DNA damage respectively (Ford et al. 2011; Namouchi et al. 2012) in Mtb. This consistency adds validity to our variant calling approach. Overall the observed diversity spared the CD4^+^ and CD8^+^ T cell epitope encoding regions of the genome providing further evidence that host adaptive immunity does not drive directional selection in Mtb genomes now at short-time scales (Coscolla et al. 2015; Comas et al. 2010; Copin et al. 2014). Diversity was concentrated in antibiotic resistance regions and strikingly also in PE/PPE genes (**Fig. 6D**) (Phelan et al. 2016). Although previous studies have generally avoided reporting short-read variant calls in PE/PPE regions, we demonstrate using read simulation, visualization of Illumina read alignments and comparison with long-read sequencing data the accuracy of the SNPs captured in our study. We found PPE genes to be more diverse in-host than PE genes and detected a signal of positive selection acting on three genes belonging to the PPE sub-family (**Fig. 7**). This indicates that PPE genes may be play an important role in the process of host-adaptation.

In addition to identifying in-host variation in 12 loci known to be involved in the acquisition of antibiotic resistance, we identified six genes and six pathways displaying diversity in-host and not known to be associated with antibiotic resistance (**Table S8, S13-14, S17**). For a subset we demonstrate similar diversity has arisen independently in separate hosts and in strains with different genetic backgrounds suggesting positive selection (**Fig. 7**). Evidence of directional selection in Mtb genomes have thus far been largely restricted to adaptation to antibiotic treatment (Didelot et al. 2016; Trauner et al. 2017; Brites and Gagneux 2015). The novel pathways showing in-host convergence may be important for interactions between host and pathogen arising from either metabolic or immune pressure. Mtb is one of a few types of bacteria that possess the capacity for *de novo* coenzyme B12/cobalamin synthesis, and this pathway has been implicated in Mtb survival in-host and Mtb growth (Rowley and Kendall 2019). We identified four genetic variants that developed in three separate patients and in three consecutive genes from the same locus *cobG*, intergenic *cobG-cobH, cobH* and *cobK* (Rv2064-Rv2067). This observation contributes to mounting evidence on the importance of this pathway for *in vivo* Mtb survival and may have implications for drug development (Minias et al. 2018; Gopinath et al. 2013). Biotin biosynthesis is also relatively unique to mycobacteria and plays an important role in Mtb growth, infection and host survival during latency (Salaemae et al. 2011). The other identified pathways include ESAT-6 protein secretion known to play a role in the modulation of host immune response by disrupting the phagosomal membrane (Clemmensen et al. 2017).

The loci found to be phylogenetically convergent and not known to be associated with antibiotic resistance, include the genes Rv0095c, *PPE18, PPE54* and *PPE60*. Although of unknown function, Rv0095c (SNP A85V) was recently associated with transmission success of an Mtb cluster in Peru (Dixit et al. 2019). Both *PPE18* and *PPE60* have been shown to interact with toll-like receptor 2 (TLR2) (Nair et al. 2009; Su et al. 2018). *PPE18* was the only gene to encode an epitope containing a SNP in-host; mutations in the epitope-encoding regions of this gene have previously been described in a set of geographically separated clinical isolates (Hebert et al. 2007). Furthermore, *PPE18* codes for one of the antigens used in the construction of the M72/AS01E vaccine candidate (Tait et al. 2019). Our results demonstrating that *PPE18* is under positive selection in the MTBC may have implications for the efficacy of this vaccine against genetically diverse Mtb strains. *PPE54* has been implicated in Mtb’s ability to arrest macrophage phagosomal maturation (phagosome-lysosome fusion) and thought to be vital for intracellular persistence (Brodin et al. 2010). The mechanism by which *PPE54* accomplishes this is unknown, but Mtb modification of phagosomal function is thought to be TLR2/TLR4-dependent (Podinovskaia et al. 2013).

Mtb is known to disrupt numerous *innate* immune mechanisms including phagosome maturation, apoptosis, autophagy as well as inhibition of MHC II expression through prolonged engagement with innate sensor toll-like receptor 2 (TLR2) among others (Ernst 2018a). SNPs in human genes involved with innate-immune pathways have been implicated in-host susceptibility to TB (Kleinnijenhuis et al. 2011; Tientcheu et al. 2017; Azad et al. 2012). Specifically, SNPs in TLR2 (thought to be the most important TLR in Mtb recognition) (Tientcheu et al. 2017) and TLR4 have been associated with susceptibility to TB disease (Kleinnijenhuis et al. 2011; Azad et al. 2012). Taken together, these observations and our results are consistent with ongoing co-evolution between humans and Mtb with evidence for reciprocal adaptive changes, leaving a signature of selection in both humans and Mtb populations (Brites and Gagneux 2015). Most co-evolution between Mtb and humans, the main reciprocal adaptations between host and pathogen are thought to have occurred long ago and as a result of long-term host-pathogen interactions (Brites and Gagneux 2015; Azad et al. 2012). Here, we observe these dynamics over the short evolutionary timescale of a single infection which has important implications for vaccine development (Brennan 2017).

## MATERIALS AND METHODS

### Sequence Data

#### Longitudinal Isolate Pairs

This study included data for 614 clinical isolates of *M. tuberculosis* that were sampled from the sputum of 307 subjects resulting in n = 307 longitudinal pairs. The sequencing data for 456 publicly available isolates was downloaded from Genbank (Benson et al. 2008), sequenced using Illumina chemistry to generate paired-end reads and came from previously published studies (T (Trauner et al. 2017), C (Casali et al. 2016), W (Walker et al. 2013), B (Bryant et al. 2013), G (Guerra-Assunção et al. 2014), X (Xu et al. 2018), H (Witney et al. 2017), P (Farhat et al. 2019)) (**Fig. S1, Table S1-S2**).

#### Replicate Isolate Pairs

This study included three types of replicate isolate pairs. (S2 - Sequenced Twice) DNA pooled from a single Mtb clinical isolate that had undergone *in vitro* expansion was sequenced in separate runs on an Illumina sequencing machine (m = 5). (C2 – Cultured & Sequenced Twice) Mtb was cultured from a single frozen clinical sample at separate time points, then sequenced on an Illumina sequencing machine after DNA extraction from culture (m = 73). (P3) Three sputum samples were obtained from a single subject within a 24 hour period (Trauner et al. 2017), cultured separately, underwent DNA extraction and then sequencing on an Illumina sequencing machine. For the purposes of this study, we compared these three isolates pairwise (m = 3).

#### Global Sequence Data

We downloaded raw sequence data for 33,873 clinical isolates from the public domain (Benson et al. 2008). Isolates had to meet the following quality control measures for inclusion in our study: (i) at least 90% of the reads had to be taxonomically classified as belonging to the *Mycobacterium tuberculosis* complex after running the trimmed FASTQ files through Kraken (Wood and Salzberg 2014) and (ii) at least 95% of bases had to have coverage of at least 10x after mapping the processed reads to the H37Rv Reference Genome.

### Epitope Collection and Analysis

CD4^+^ T and CD8^+^ T cell epitope sequences were downloaded from the Immune Epitope Database (Vita et al. 2014) on May 23rd, 2018 according to criteria described previously (Coscolla et al. 2015) [linear peptides, *M. tuberculosis* complex (ID:77643, Mycobacterium complex), positive assays only, T cell assays, any MHC restriction, host: humans, any diseases, any reference type] yielding a set of 2,031 epitope sequences (**Table S11**). We mapped each epitope sequence to the genes encoded by the H37Rv Reference Genome (Cole et al. 1998) using BlastP with an e-value cutoff of 0.01 (**Fig. S3A**). We retained only epitope sequences that mapped to at least 1 region in H37Rv (due to sequence homology, some epitopes mapped to multiple regions) and whose BlastP peptide start/end coordinates matched those specified in IEDB (n = 1,949 representing 1,505 separate epitope entries in IEDB). We then filtered out any epitopes occurring in Mobile Genetic Elements which resulted in a final set of 1,875 epitope sequences, representing 348 genes (antigens) used for downstream analysis. The distribution of peptide lengths for this final set of epitopes is given in **Fig. S3B**. Since many of these epitope sequences overlap, we constructed non-redundant epitope concatenate sequences for each antigen (n = 348) gene (Coscolla et al. 2015; Stucki et al. 2016; Comas et al. 2010). The regions of each antigen not encoding an epitope were concatenated into a non-epitope sequence for that gene.

### Gene Sets

Every gene on H37Rv was classified into one of six non-redundant gene categories according to the following criteria: (i) genes identified as belonging to the PE/PPE family of genes unique to pathogenic mycobacteria, though to influence immunopathogenicity and characterized by conserved proline-glutamate (PE) and proline-proline-glutamate (PPE) motifs at the N protein termini (Brennan and Delogu 2002; Comas et al. 2010; Phelan et al. 2016) were classified as *PE/PPE* (n = 167), (ii) genes flagged as being associated with antibiotic resistance (Farhat et al. 2013) were classified into the *Antibiotic Resistance* category (n = 28), (iii) genes encoding a CD4^+^ or CD8^+^ T-cell epitope (Coscolla et al. 2015; Comas et al. 2010) (but not already classified as a PE/PPE or Antibiotic Resistance gene) were classified as an *Antigen* (n = 257), (iv) genes required for growth *in vitro* (Sassetti et al. 2003) and *in vivo* (Sassetti and Rubin 2003) and not already placed into a category above were classified as *Essential* genes (n = 682), (v) genes flagged as transposases, integrases, phages or insertion sequences were classified as *Mobile Genetic Elements* (Comas et al. 2010) (n = 108), (vi) any remaining genes not already classified above were placed into the *Non-Essential* category (n = 2752) (**Table S4**).

### Illumina Sequencing FastQ Processing and Mapping to H37Rv

The raw sequence reads from all sequenced isolates were trimmed with Prinseq (Schmieder and Edwards 2011) (settings: -min_qual_mean 20) (version 0.20.4) then aligned to the H37Rv Reference Genome (Genbank accession: NC_000962) with the BWA mem (Li and Durbin 2009) algorithm (settings: -M) (version 0.7.15). The resulting SAM files were then sorted (settings: SORT_ORDER = coordinate), converted to BAM format and processed for duplicate removal with Picard (http://broadinstitute.github.io/picard/) (version 2.8.0) (settings: REMOVE_DUPLICATES = true, ASSUME_SORT_ORDER = coordinate). The processed BAM files were then indexed with Samtools (Li et al. 2009). We used Pilon (Walker et al. 2014) on the resulting BAM files to call bases for all reference positions corresponding to H37Rv from pileup (settings: --variant).

### Empirical Score for Difficult-to-Call Regions

We extracted DNA from 15 Mtb isolates (Epperson and Strong 2020), for which we had Illumina sequencing reads, to undergo PacBio sequencing (**Supplementary Note 2)**. Together, with public PacBio and Illumina sequencing data (Chiner-Oms et al. 2019), we compiled 31 pairs of sequencing reads for comparison of variant calling between PacBio long-reads and Illumina short-reads (**Table S20**). Using the 31 isolates for which both Illumina and a complete PacBio assembly were available (**Supplementary Note 2)**, we evaluated the empirical base-pair recall (EBR) of all base-pair positions of the H37rv reference genome (Marin et al. 2020). For each sample, the alignments (from minimap2) of each high confidence genome assembly to the H37Rv genome were used to infer the true nucleotide identity of each base pair position. To calculate the empirical base-pair recall, we calculated what % of the time our Illumina based variant calling pipeline, across 31 samples, confidently called the true nucleotide identity at a given genomic position. If Pilon variant calls did not produce a confident base call *(Pass)* for the position, it did not count as a correct base call. This yields a metric ranging from 0.0 – 1.0 for the consistency by which each base-pair is both confidently and correctly sequenced by our Illumina WGS based variant calling pipeline for each position on the H37Rv reference genome. An H37Rv position with an EBR score of x% indicates that the base calls made from Illumina sequencing and mapping to H37Rv agreed with the base calls made from the PacBio *de novo* assemblies (**Supplementary Note 2**) in x% of the Illumina-PacBio pairs. We masked difficult-to-call regions by dropping H37Rv positions with an EBR score below 0.8 as part of our variant calling procedure.

### Variant Calling

#### Single Nucleotide Polymorphism (SNP) Calling

To prune out low-quality base calls that may have arisen due to sequencing or mapping error, we dropped any base calls that did not meet any of the following criteria (Copin et al. 2016): (i) the call was flagged as either *Pass* or *Ambiguous* by Pilon, (ii) the reads aligning to that position supported at most 2 alleles (ensuring that 1 allele matched the reference allele if there were 2), (iii) the mean base quality at the locus was > 20, (iv) the mean mapping quality at the locus was > 30, (v) none of the reads aligning to the locus supported an insertion or deletion, (vi) a minimum coverage of 25 reads at the position, (vii) the EBR score for the position ≥0.80, and (viii) the position is not located in a mobile genetic element region of the reference genome. We then used the Pilon-generated (Walker et al. 2014) VCF files to calculate the frequencies for both the reference and alternate alleles, using the *INFO.QP* field (which gives the proportion of reads supporting each base weighted by the base and mapping quality of the reads, *BQ* and *MQ* respectively, at the specific position) to determine the proportion of reads supporting each base for each locus of interest.

#### Additional SNP Filtering for Isolate Pairs

To call SNPs (and corresponding changes in allele frequencies) between pairs of isolates (Replicate and Longitudinal pairs), we required: (i) *SNP Calling* filters be met, (ii) the number of reads aligning to the position is below the 99th percentile for all of the calls made for that isolate, (iii) the call at that position passes all filters for each isolate in the pair, and (iv) SNPs in *glpK* were dropped as mutants arising in this gene are thought to be an artifact of *in vitro* expansion (Pethe et al. 2010; Vargas and Farhat 2020); we detected four non-synonymous SNPs in *glpK* (ΔAF ≥ 25%) between three longitudinal pairs (mean ΔAF=64%) and five non-synonymous SNPs in *glpK* (ΔAF ≥ 25%) among five replicate pairs (mean ΔAF=45%).

#### Additional SNP Filtering for Antibiotic Resistance Loci Analysis

To call SNPs (and corresponding minor changes in allele frequencies) between pairs of isolates (Longitudinal Pairs), we required: (i) *SNP Calling* filters be met, (ii) *Additional SNP Filtering for Isolate Pairs* filters be met, (iii) 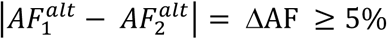, (iv) if 5% ≤ ΔAF < 20%, then the SNP was only retained if each allele (across both isolates) with AF > 0% was supported by at least 5 reads (ensuring that at least 5 reads supported each minor allele at lower values of ΔAF), (v) the SNP was classified as either intergenic or non-synonymous, (vi) the SNP was located in a gene, intergenic region or rRNA coding region associated with antibiotic resistance (**Table S5**).

#### Additional SNP Filtering for Heterogenous SNPs

To call heterogenous SNPs in each isolate for a pair of longitudinal isolates, we required: (i) *SNP Calling* filters be met, (ii) the number of reads aligning to the position is below the 99th percentile for all of the calls made for that isolate, (iii) the alternate allele frequency for the SNP is such that 25% ≤ *AF*^*alt*^ ≤ 75%, (iv) the position is not located in a mobile genetic element region of the reference genome, (v) the position is not located in a PE/PPE region of the reference genome.

#### Additional SNP Filtering for Global Isolates

To call alleles in our global and genetically diverse set of isolates, we required: (i) *SNP Calling* filters be met, with the modifications that (ii) the call was flagged as *Pass* by Pilon and (iii) 1 allele (either the reference or an alternate) was supported by at least 90% of the reads regardless of whether the other ≤ 10% of reads supported 1, 2, or 3 alleles.

### Mixed Lineage and Contamination Detection for Longitudinal and Replicate Isolate Pairs

#### Kraken

To filter out samples that may have been contaminated by foreign DNA during sample preparation, we ran the trimmed reads for each longitudinal and replicate isolate through Kraken2 (Wood and Salzberg 2014) against a database (Goig et al. 2020) containing all of the sequences of bacteria, archaea, virus, protozoa, plasmids and fungi in RefSeq (release 90) and the human genome (GRCh38). We calculated the proportion reads that were taxonomically classified under the *Mycobacterium tuberculosis* Complex (MTBC) for each isolate and implemented a threshold of 95%. An isolate pair was dropped if either isolate had less than 95% of reads aligning to MTBC.

#### F2

To further reduce the effects of contamination, we aimed to identify samples that may have been subject to inter-lineage mixture samples resulting from of a co-infection (F2). We computed the F2 lineage-mixture metric for each longitudinal and replicate isolate (**Fig. 1B**). We wrote a custom script to carry out the same protocol for computing F2 as previously described (Wyllie et al. 2018). Briefly, the method involves calculating the minor allele frequencies at lineage-defining SNPs (Coll et al. 2014). From 64 sets of SNPs that define the deep branches of the MTBC (Coll et al. 2014), we considered the 57 sets that contain more than 20 SNPs to obtain better estimates of minor variation (Wyllie et al. 2018; Coll et al. 2014). For each SNP set *i*, (i) we summed the total depth and (ii) the number of reads supporting the most abundant base (at each position) over all of the reference positions (SNPs) that met our mapping quality, base quality and insertion/deletion filters, which yields *d*_*i*_ and *x*_*i*_ respectively. Subtracting these two quantities yields the minor depth for SNP set *i, m*_*i*_ = *d*_*i*_ − *x*_*i*_. The minor allele frequency estimate for SNP set *i* is then defined as *p*_*i*_ = *m*_*i*_ / *d*_*i*_. Doing this for all 57 SNP sets gives {*p*_1_, *p*_2_, …, *p*_57_}. We then sorted {*p*_1_, *p*_2_, …, *p*_57_} in descending order and estimated the minor variant frequency for all of the reference positions (SNPs) corresponding to the top 2 sets (highest *p*_*i*_ values) which yields the F2 metric. Letting *n*2 be the number of SNPs in the top 2 sets, then 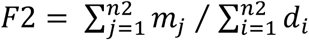. Isolate pairs were dropped if the F2 metric for either isolate passed the F2 threshold set for mixed lineage detection (**Fig. 1B** and **Fig. S1**).

### Pre-existing Genotypic Resistance

We determined pre-existing resistance for a subject (with a pair of longitudinal isolates) by scanning the first isolate for the detection of at least 1 of 177 SNPs predictive of resistance with AF ≥ 75% (from a minimal set of 238 variants (Farhat et al. 2016)). We excluded predictive indels and the *gid* E92D variant as the latter is likely a lineage marking variant that is not indicative of antibiotic resistance. We defined pre-existing multidrug resistance for a subject by scanning the first isolate collected for detection of at least 1 SNP predictive of Rifampicin resistance (14/178 predictive SNPs) and at least 1 SNP predictive of Isoniazid resistance (18/178 predictive SNPs).

### True & False Positive Rate Analysis for Heteroresistant Mutations

To determine the predictive value of low-frequency heteroresistant alleles, we classified SNPs as fixed if the alternate allele frequency in the 2^nd^ isolate collected from the subject was at least 75% (alt AF_2_ ≥ 75%). We first dropped SNPs for which alt AF_1_ ≥ 75% and alt AF_2_ ≥ 75% (high frequency mutant alleles in both isolates). We then set a threshold *(F*_*i*_) for the alternate allele frequency detected in the 1^st^ isolate collected from the subject (alt AF_1_) and predicted whether an alternate allele would rise to a substantial proportion of the sample (alt AF_2_ ≥ 75%) as follows:

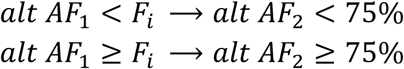

We classified every SNP as True Positive (TP), False Positive (FP), True Negative (TN) or False Negative (FN) according to:

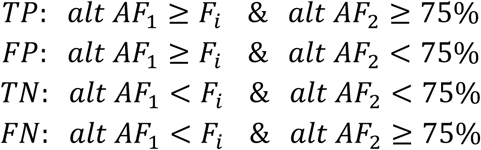

True Positive Rates (TPR) and False Positive Rates (FPR) were calculated as:

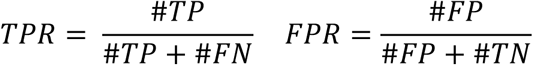

Finally, we made predictions for all SNPs and calculated the TPR and FPR for all values of *F*_*i*_ ∈ {0%, 1%, 2%, …, 98%, 99%, 100%}.

### Mutation Density Test

The method to detect significant variation for a given locus amongst pairs of sequenced isolates has been described previously (Farhat et al. 2014). Briefly, let 𝒩_*j*_ ∼ *Pois*(*λ*_*j*_) be a random variable for the number of SNPs detected across all isolate pairs (for the in-host analysis this is the collection of longitudinal isolate pairs for all subjects) for gene *j*. Let (i) *N*_*i*_ = number of SNPs across all pairs for gene *i*, (ii) |*g*_*i*_| = length of gene *i*, (iii) *P* = number of genome pairs and (iv) *G* = the number of genes across the genome being analyzed (all genes in the essential, non-essential, antigen, antibiotic resistant and family protein categories).

Then the length of the genome (concatenate of all genes being analyzed) is given by 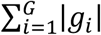 and the number of SNPs across all genes and genome pairs is given by 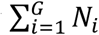. The null rate for 𝒩_*j*_ is given by the mean SNP distance between all pairs of isolates, weighted by the length of gene *j* as a fraction of the genome concatenate and number of isolate pairs:

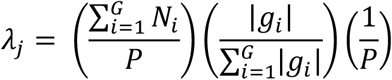

The p-value for gene *j* is then calculated as Pr (*N*_*i*_ > 𝒩_*j*_). We tested 3,386 genes for mutational density and applied Bonferroni correction to determine a significance threshold. We determine a gene to have a significant amount of variation if the assigned 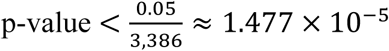.

### Nucleotide Diversity

We define the nucleotide diversity (*π*_*g*_)for a given gene *g* as follows: (i) let |*gene*_*g*_| = base-pair length of the gene, (ii) *N*_*i,j*_ = number of in-host SNPs (independent of the change in allele frequency for each SNP) between the longitudinal isolates for subject *i* occurring on gene *j* and (iii) *P* = number of subjects. Then

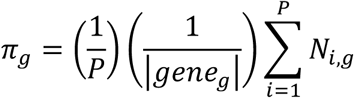

Correspondingly, let *G* be a category consisting of *M* genes, then the average nucleotide diversity for *G* is given by:

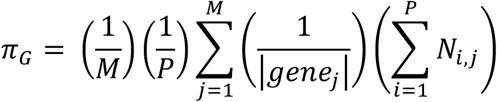

### SNP calling simulations in repetitive genomic regions

Certain repetitive regions of the *Mycobacterium tuberculosis* genome (ESX, PE/PPE loci) may give rise to false positive and false negative variant calls due to the mis-alignment of short-read sequencing data. To test the rate of false negative and false positive SNP calls in genes with *in-host* SNPs (**Fig. 5, Table S8**) we collected the set of non-redundant SNPs observed in these loci (**Table S10**). Next, we collected a set of publicly available reference genomes (**Table S9**) and introduced these mutations into the respective loci positions in the reference genomes. We then simulated short-read Illumina sequencing data of comparable quality to our sequencing data from these altered reference genomes. Using our variant-calling pipeline to call polymorphisms, we then estimated the number of true and false positive SNP calls for each gene, based off of how many introduced SNPs were called (true positives), how many introduced SNPs were not called (false negatives) and how many spurious SNPs were called (false positives). A schematic of our simulation methodology is given in **Fig. S5**, a detailed explanation is given in **Supplementary Note 1** and the results of our simulations (given in **Fig. S6)** confirm a low false-positive rate.

### Global Lineage Typing

We determined the global lineage of each longitudinal (*N* = 614) and global isolate (*N* = 32,033) using base calls from Pilon-generated VCF files and a 62-SNP lineage-defining diagnostic barcode from a previously published study (Coll et al. 2014).

### Phylogenetic Convergence Analysis & t-SNE Visualization

#### Construction of Genotypes Matrix

We detected SNP sites at 878,244 H37Rv reference positions (of which 61,918 SNPs were not biallelic) among our global sample of 33,873 isolates. After excluding SNP sites with rare minor alleles (sites in which alternate alleles were called in < 5 isolates) we retained SNPs at 146,874 positions. We constructed a 146,874 x 33,873 genotypes matrix (coded as 0:A, 1:C, 2:G, 3:T, 9:Missing) and filled in the matrix for the allele supported at each SNP site for each isolate according to the *SNP Calling* filters outlined above. If a base call at a specific reference position for an isolate did not meet the filter criteria that allele was coded as *Missing*.

Excluding 2,348 SNP sites that had an EBR score < 0.80, another 1,509 SNP sites located within mobile genetic element regions, and 3,220 SNP sites in with missing calls in > 25% of isolates yielded a genotypes matrix with dimensions 139,797 × 33,873. Next, we excluded 1,518 isolates with missing calls in > 25% of SNP sites yielding a genotypes matrix with dimensions 139,797 × 32,355. We used a previously published 62-SNP barcode (Coll et al. 2014) to type the global lineage of each isolate in our sample. We further excluded 322 isolates that did not get assigned a global lineage, another 100 isolates that were used in our longitudinal analysis (**Table S3**), 152 isolates typed as *Mycobacterium bovis*, and 35 isolates typed as lineage 7. To improve computational efficiency and runtime, we randomly down sampled (using Python’s random.sample() function) our collection of lineage 2 and lineage 4 isolates to include 7,000 isolates of each lineage in our sample. This excluded 1,064 lineage 2 isolates and 10,330 lineage 4 isolates. These steps yielded a genotypes matrix with dimensions 139,797 × 20,352. Finally, we excluded 10,899 SNP sites from this filtered genotypes matrix in which the minor allele count = 0. The genotypes matrix used for downstream analysis had dimensions 128,898 x 20,352 representing 128,898 SNP sites across 20,352 isolates. The global lineage breakdown of the 20,352 isolates was: L1=2,802, L2=7,000, L3=3,352, L4=7,000, L5=101, L6= 97.

#### Phylogenetic Convergence Test

We tested 141/174 in-host SNPs in which the alternate (mutant) allele frequency increased substantially across sampling (**Table S8**) for phylogenetic convergence. We scanned a set 20,352 global and genetically diverse isolates from our 128,898 × 20,352 genotypes matrix for these SNPs (**Table S18**). To determine phylogenetic convergence for a given SNP site, we required that the alternate allele (called against the H37Rv reference) be present but not fixed in at least three global lineages. More specifically, we required that the alternate allele be detected in at least 1 isolate within each lineage and not in > 95% of the isolates in that lineage. Twenty-six SNP sites across 12 genes and 3 intergenic regions were detected as having a signal of phylogenetic convergence (**Table S19**).

#### t-SNE Visualization

To construct the t-SNE plots that captured the genetic relatedness of the 20,352 isolates in our sample, we first constructed a pairwise SNP distance matrix. To efficiently compute this using our 128,898 x 20,352 genotypes matrix, we binarized the genotypes matrix and used sparse matrix multiplication implemented in Scipy to compute five 20,352 × 20,352 similarity matrices (Virtanen et al. 2020). We constructed a similarity matrix for each nucleotide (***A, C, G, T***) where row *i*, column *j* of the similarity matrix for nucleotide *x* stored the number of *x*’s that isolate *i* and isolate *j* shared in common across all SNP sites. The fifth similarity matrix (***N***) stored the number of SNP sites in which neither isolate *i* and isolate *j* had a missing value. The pairwise SNP distance matrix (***D***) was then computed as ***D*** = ***N*** − (***A* + *C* + *G* + *T***). ***D*** had dimensions 20,352 × 20,352 where row *i*, column *j* stored the number of SNP sites in which isolate *i* and isolate *j* disagreed. We used ***D*** as input into a t-SNE algorithm implemented in Scikit-learn (Pedregosa et al. 2011) (settings: perplexity = 40, n_components = 2, metric = “precomputed”, n_iter = 1000, learning_rate = 500) to compute the embeddings for all 20,352 isolates in our sample. We used these embeddings to visualize the genetic relatedness of the isolates in two dimensions and colored isolates (points on the t-SNE plot) by lineage (**Fig. 7A**). For visualizing specific mutations, isolates were colored according to whether or not the alternate (mutant) allele was called (**Fig. 7B-F, S10-S11**).

### Data Analysis and Variant Annotation

Data analysis was performed using custom scripts run in Python and interfaced with iPython (Pérez and Granger 2007). Statistical tests were run with Statsmodels (Seabold and Perktold 2010) and Figures were plotted using Matplotlib (Hunter 2007). Numpy (Van Der Walt et al. 2011), Biopython (Cock et al. 2009) and Pandas (McKinney and others 2010) were all used extensively in data cleaning and manipulation. Functional annotation of SNPs was done in Biopython (Cock et al. 2009) using the H37Rv reference genome and the corresponding genome annotation. For every SNP called, we used the H37Rv reference position provided by Pilon (Walker et al. 2014) generated VCF file to extract any overlapping CDS region and annotated SNPs accordingly. Each overlapping CDS regions was then translated into its corresponding peptide sequence with both the reference and alternate allele. SNPs in which the peptide sequences did not differ between alleles were labeled *synonymous*, SNPs in which the peptide sequences did differ were labeled *non-synonymous* and if there were no overlapping CDS regions for that reference position, then the SNP was labeled *intergenic*.

### Pathway Definitions

We used SEED (Overbeek et al. 2013) subsystem annotation to conduct pathway analysis and downloaded the subsystem classification for all features of *Mycobacterium tuberculosis* H37Rv (id: 83332.1) (**Table S15**). We mapped all of the annotated features from SEED to the annotation for H37Rv. Due to the slight inconsistency between the start and end chromosomal coordinates for features from SEED and our H37Rv annotation, we assigned a locus from H37Rv to a subsystem if both the start and end coordinates for this locus fell within a 20 base-pair window of the start and end coordinates for a feature in the SEED annotation (**Table S16**).

## ACKNOWLEDGEMENTS

We thank the members of the Farhat lab for helpful discussions and comments on the research project and manuscript. We thank S. Fortune, N. Hicks & D. Warner for helpful suggestions on the manuscript. We thank A. Narayan for helpful suggestions on constructing t-SNE visualizations for phylogenetic convergence. R.V.J. was supported by the National Science Foundation Graduate Research Fellowship under Grant No. DGE1745303. Portions of this research were conducted on the O2 High Performance Compute Cluster, supported by the Research Computing Group, at Harvard Medical School.

## AUTHOR CONTRIBUTIONS

R.V.J. and M.R.F. conceived, designed and conducted the study. R.V.J. and M.R.F. drafted the manuscript with input from all authors. L.F. and M.M. provided bioinformatics support. L.E.E., D.D., M. Salfinger and M. Strong cultured Mtb isolates and performed DNA extraction in preparation for PacBio sequencing. M.Smith and I.O. prepared libraries and performed PacBio sequencing runs.

## DATA AND MATERIALS AVAILABILITY

All Mtb sequencing data was collected from previously published studies and is publicly available. Individual accession numbers for the Mtb genomes analyzed in this study can be found in **Table S2** and information on which studies from which the data was generated can be found in the **Methods, Fig. S1** and **Table S1**. All packages and software used in this study have been noted in the **Methods**. Custom scripts written in python version 2.7.15 were used to conduct all analyses and interfaced via Jupyter Notebooks. All scripts and notebooks will be uploaded to a GitHub repository upon acceptance of this manuscript for publication.

