## Supplementary Text and Figures for "In-host population dynamics of *M. tuberculosis* during treatment failure"

### TABLE OF CONTENTS


- Supplementary Note 1 – SNP Calling Simulations
  - Reference Genome Collection
  - Mapping CDS regions from Reference Genomes to H37Rv
  - Filtering Low-Quality Mapped Reference Genomes
  - Altering RefGenomes at SNP Test Sites
  - Simulating Reads from Complete Genomes
  - Mapping Simulated Reads to H37Rv and Calling SNPs
  - Calling SNPs with MUMmer
  - True & False Positive SNP Call Analysis
- Supplementary Note 2 – PacBio Assembly vs. Illumina Mapping SNP Calling
  - DNA extraction and PacBio Sequencing of Mtb Isolates
  - PacBio *de novo* Assembly, Genome Polishing, and Variant Calling
- Supplementary Figures 1-11
- Supplementary Table Descriptions 1-20
- References for Supplementary Information

### SUPPLEMENTARY NOTE 1 – SNP Calling Simulations

#### Reference Genome Collection

We downloaded 60 reference genomes (RefGenome) (i.e. completely assembled *Mycobacterium tuberculosis* genomes) from NCBI (Genbank accession IDs can be found in **Table S9**). We limited our collection to genomes for which there were corresponding annotation files.

#### Mapping CDS regions from Reference Genomes to H37Rv

Since the regions of interest were repetitive loci that have many homologies elsewhere in the genome, we were unable to use traditional alignment methods to map the genes of interest from H37Rv to the other RefGenomes. Instead, we made use of the clonal structure of the Mtb genome to construct gene mappings from H37Rv to the RefGenomes as follows (**Fig. S5A**):

1. For each gene $g$ annotated in H37Rv, collect the set of gene lengths 5 genes upstream and 5 genes downstream of $g$ from H37Rv. Compare the set of 11 H37Rv gene lengths to every set of 11 consecutive gene neighborhoods on the RefGenome and assign a score based off of the intersection of each pair of sets.
2. Look at the gene neighborhood(s) with the top score after scanning the RefGenome and pairwise globally align (Cock et al. 2009) $g$ to every gene in the top scoring neighborhood using the following criteria: (i) identical characters are given 2 points, (ii) 1 point is deducted for each non-identical character, (iii) 2 points are deducted for opening a gap, (iv) 2 points are deducted for extending a gap.
3. Take the top scoring alignment $r$ and assign a mapping from H37Rv gene $g$ to RefGenome gene $r$ if (i) the pairwise alignment score is $> 0$ and (ii) the base pair length of $g$ and $r$ are equivalent (the latter ensures correct placement of mutations in

downstream analysis). If either of these criteria is not met, then we do not assign a mapping from $g$ to any CDS region on that RefGenome.

**Filtering Low-Quality Mapped Reference Genomes**

To assess the quality of the mappings from H37Rv to the set of RefGenomes, we compared the reference position start coordinates of each assigned mapping between each RefGenome and H37Rv. Again, making use of Mtb clonality, we reasoned that the genomic structure of each pair of genomes is similar (if each RefGenome is indexed to start at the first gene on H37Rv *Rv0001*, then well mapped RefGenomes will have mapped genes that are located within a neighborhood of the coordinates from H37Rv). To test this (for each RefGenome), we took the absolute difference between the start coordinates for all of the mapped genes between the RefGenome and H37Rv. We then averaged these differences across all gene mappings between both genomes. This measures the conservation (of the ordering) of the mapped genes between each pair of genomes (H37Rv & RefGenome) and gives an indication of how successful the mappings were on a global scale. We downloaded and mapped genes for 60 Genome Assemblies from GenBank (Benson et al. 2008) and assessed the quality of each set of mappings using the measure described above (**Fig. S5B-C**). We excluded 6 RefGenomes on the basis of sporadic gene mappings against H37Rv which was determined by looking at the distribution of the mapping measure for all 60 assemblies. We kept the remaining 54 genomes for use in the simulations (**Table S9**).

**Altering RefGenomes at SNP Test Sites**

We make use of the set of the (non-redundant) observed in-host SNPs across all genes (**Fig. 5D, Table S10**). We alter each RefGenome by introducing mutations (that correspond to the aforementioned SNPs) into the genes successfully mapped to H37Rv, ensuring that the new bases differ from the corresponding base positions on H37Rv. Since successful mappings require that the mapped genes be the same length, the mutations are introduced into the same site on the RefGenome with respect to the gene specific coordinates (i.e. a gene $n$ bp long will have coordinates $\{1, 2, \cdots, n-1, n\}$ from $5' \rightarrow 3'$). We store information pertaining to which bases were altered for each RefGenome $\{SNP\ set\ \boldsymbol{\beta}\}$. No simulations are run for genes on RefGenomes that are not successfully mapped to H37Rv.

**Simulating Reads from Complete Genomes**

To validate our SNP calling methodology using the set of RefGenomes, we used ART (Huang et al. 2011) to simulate short-read sequencing data altered versions of the RefGenomes (**Fig. S5B**). Since the aim of our simulations was to study the quality of our variant calls on our real data, we simulated data for each (altered) RefGenome that was of comparable quality to our real sequencing data: Illumina HiSeq 1000, read length of 100bp, mean coverage of 80x, paired end reads, 200bp mean size of DNA fragments, 25bp standard deviation of DNA fragment size (settings: -ss HS10 -l 100 -f 80 -p -m 200 -s 25).

**Mapping Simulated Reads to H37Rv and Calling SNPs**

Next we mapped the pool of simulated reads from the altered RefGenomes against the H37Rv reference genome and called SNPs according to most of the same procedures and WGS filters outlined in **Methods**. However, in this instance we called SNPs at reference positions that supported an alternate allele and required that calls were flagged as *Pass* by Pilon (where the alternate allele frequency was $\geq 75\%$ and no *Ambiguous*, *Low Coverage*, or *Deletion* flags were present at that position). For each RefGenome, this yielded the set of SNPs (between the altered RefGenome and H37Rv) called by our pipeline $\{SNP\ set\ \mathbf{B}\}$ (**Fig. S5B**).

**Calling SNPs with MUMmer**

We used Mummer3 (Kurtz et al. 2004) to call SNPs between H37Rv and each (unaltered) RefGenome. We aligned each pair of genomes and called SNPs between the alignments using the following commands:

1) nucmer -mum H37Rv.fasta RefGenome.fasta
2) delta-filter -r -q H37Rv_RefGenome.delta > H37Rv_RefGenome.filter
3) show-snps -Clr -T H37Rv_RefGenome.filter > H37Rv_RefGenome.snps

The resulting SNP calls yielded the set of SNPs between each of the unmodified (unaltered) RefGenomes and H37Rv $\{SNP\ set\ \mathbf{A}\}$ (**Fig. S5B**).

**True & False Positive SNP Call Analysis**

To calculate the number of *true positives* and *false positives* with regard to our SNP calling pipeline for each gene $g$ of interest (**Fig. S6**), we define the following sets of H37Rv coordinates for each RefGenome:

- $\boldsymbol{\beta}$ - SNPs introduced into (altered) RefGenome
- $\boldsymbol{A}$ - SNPs called between (unaltered) RefGenome & H37Rv
- $\boldsymbol{B}$ - SNPs called between (altered) RefGenome & H37Rv
- $\boldsymbol{C}$ - all reference positions (or coordinates) on H37Rv

The set of coordinates where an alternate allele was introduced into the RefGenome and called by the pipeline (true positive SNPs for gene $g$) is given by:

$$TP_g = (B_g \setminus A_g) \cap (\beta_g \setminus A_g)$$

where we normalize by SNP set $A_g$ to make sure we're only accounting for test SNPs in our computations. The set of coordinates where an alternate allele was note introduced and called by the pipeline (false positive SNPs for gene $g$) is given by:

$$FP_g = \left((B_g \setminus A_g) \cap C_g\right) \setminus TP_g$$

The set of coordinates where an alternate allele was introduced but was not called by the pipeline (false negative SNPs for gene $g$) is given by:

$$FN_g = (\beta_g \setminus A_g) \setminus TP_g$$

The results of our simulations (**Fig. S6**) indicate that the number of true positive calls is consistent with the number of known SNPs across all genes and simulations. Perhaps more importantly, our results also suggest that false positive calls are rarely made for any SNP in our sample. Thus, while we may not have called all of the existing variation between paired isolates (false negative calls), it is unlikely that we called non-existing variation between any pair of isolates (false positives). That is, false-positive SNPs are rarely called, even in repetitive loci such as the PE/PPE gene family, supporting our decision to keep all SNP calls for downstream analysis.

### SUPPLEMENTARY NOTE 2 – PacBio Assembly vs. Illumina Mapping SNP Calling

#### DNA extraction and PacBio Sequencing of Mtb Isolates

DNA extraction was performed according to a published protocol (Epperson and Strong 2020). Approximately 1 mg of high molecular weight genomic DNA was used as input for SMRTbell preparation, according to the manufacturer's specifications (SMRTbell Template Preparation Kit 1.0, Pacific Biosciences, https://www.pacb.com/wp-content/uploads/2015/09/Procedure-Checklist-20-kb-Template-Preparation-Using-BluePippin-Size-Selection.pdf). Briefly, HMW gDNA was sheared to 20kb using the Covaris g-tube at 4500 rpm. Following shearing, gDNA underwent DNA damage repair, ligation to SMRTbell adaptors and exonuclease treatment to remove any unligated gDNA. At least 500 ng final SMRTbell library per sample was cleaned with AMPure PB beads and 3-50 kb fragments were size selected using the BluePippin system on 0.75% agarose cassettes and S1 ladder, as specified by the manufacturer (Sage Science). Size selected SMRTbell libraries were annealed to sequencing primer and bound to the P6 polymerase prior to loading on the RSII sequencing system (Pacific Biosciences). Sequencing was performed using C4 chemistry and 240-minute movies. Following data collection, raw data was converted into subreads for subsequent analysis using the RS_Subreads.1 pipeline within SMRTPortal (version 2.3), the web-based bioinformatics suite for analysis of RSII data.

#### PacBio *de novo* Assembly, Genome Polishing, and Variant Calling

PacBio and Illumina sequencing data was available for 34 clinical Mtb isolates (Marin et al. 2020; Chiner-Oms et al. 2019). We used Flye (Kolmogorov et al. 2019) to *de novo* assemble the raw PacBio subreads from these 34 isolates (settings: --pacbio-raw --genome-size 5m) (version 2.5). If Flye identified the presence of a circular contig, Circlator (Hunt et al. 2015) was used to set the start each assembly at the DnaA locus. PacBio's bax2bam function (settings: --subread) was used to convert PacBio legacy BAX files to BAM format. We ran PacBio's implementation of Minimap2 (Li 2018) (pbmm2) to map and sort raw PacBio subreads to the *de novo* assembly. We iteratively polished the assembly three times by running the Quiver algorithm (Chin et al. 2013) and used Samtools (Li et al. 2009) to index the fasta files from the resulting assemblies. Thirty-one of our 34 samples assembled into a single circular contig (**Table S20**). We excluded three isolates that did not have a single circular assembly from downstream analysis. To call SNPs relative to the H37Rv reference, we used Minimap2 (Li 2018) to align each PacBio assembly to the H37Rv reference sequence. We used the *paftools.js call* utility included with Minimap2 to generate variant calls from each assembly to reference alignment.

### SUPPLEMENTARY FIGURES

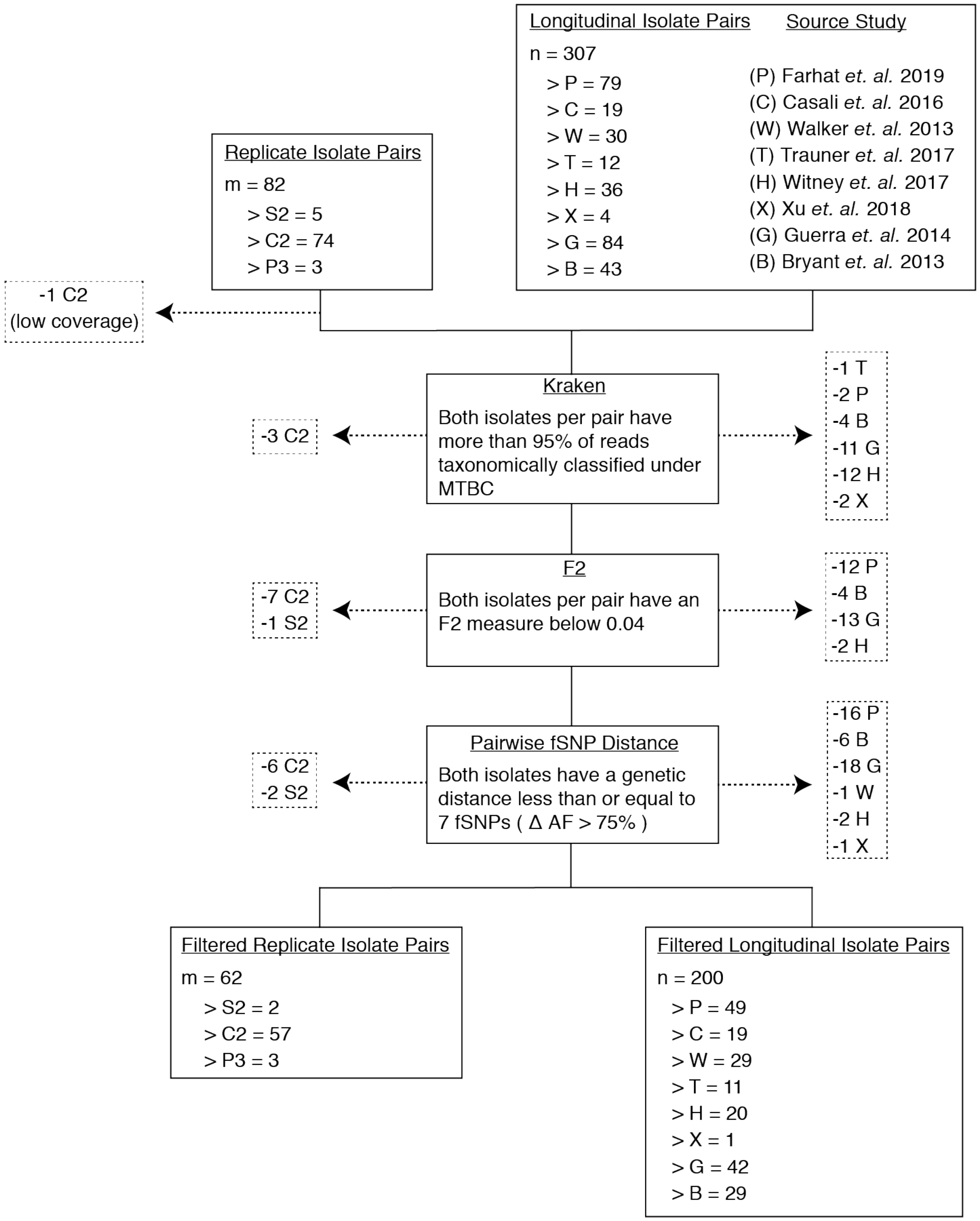


**Fig. S1. Filtering out laboratory-contaminated samples and subjects with mixed infections.** We implemented several filters to mitigate the effects of contamination from laboratory error or samples from co-infected hosts (**Methods**). Our analysis included three types of replicate pairs

(S2, C2, P3) and longitudinal pairs from eight studies (P, C, W, T, B, G, X, H) (**Methods**). At each step, we filtered out any pair of isolates if at least one isolate failed to pass the filter in place (indicated by dashed arrows). First, we used Kraken to filter out isolates that had less than 95% of reads taxonomically classified under MTBC. Second, we filtered out isolates that did not meet the F2 threshold. Third, we filtered out isolate pairs that had a genetic distance greater than 7 fixed SNPs. Our final filtered isolate pair sets included 62 replicate isolate pairs and 200 longitudinal isolate pairs.

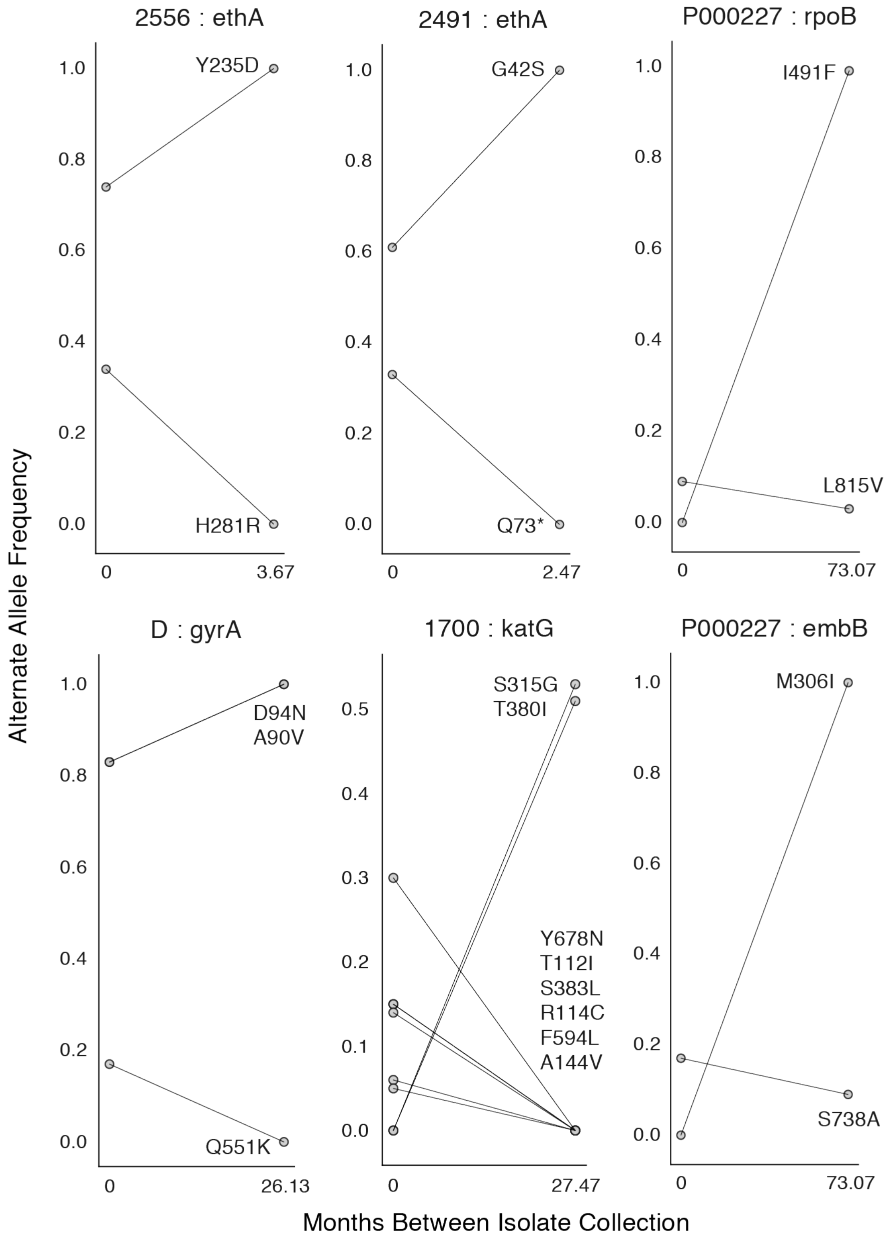


**Fig. S2. Mutant allele trajectories consistent with clonal interference.** Several examples of co-occurring mutant alleles and their allele frequency trajectories between longitudinal isolate collection demonstrate genetic diversity patterns consistent with competing clones in-host. Each mutant allele is labeled with amino acid encoded by the reference allele, H37Rv codon position, and amino acid encoded by the mutant allele.

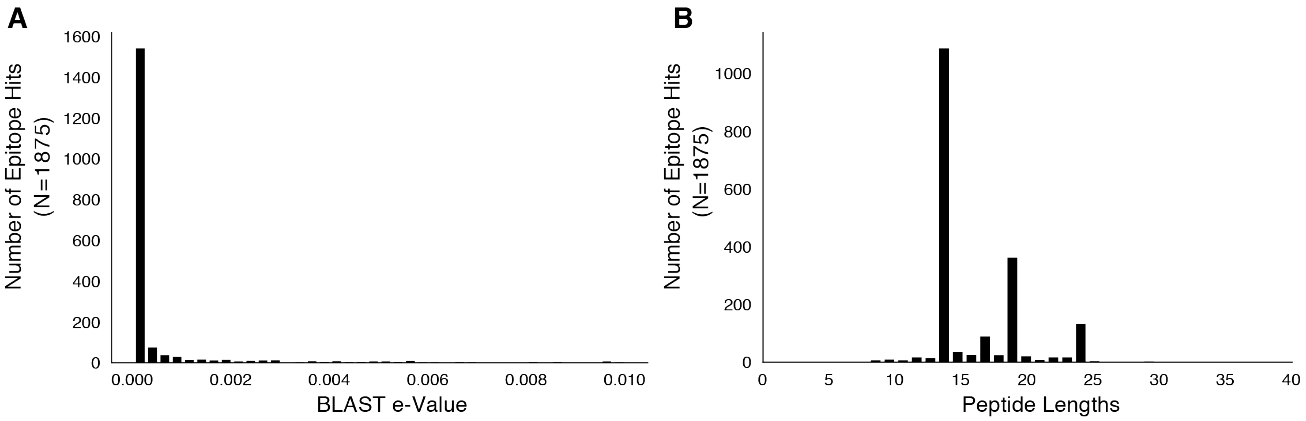


**Fig. S3. Basic characteristics of epitopes used in analysis.** We downloaded a set of 2,031 epitope peptide sequences from IEDB (Vita et al. 2014) and used BLASTP to map these peptide sequences to H37Rv imposing an e-value cut-off of 0.01 (**Methods**). (**A**) The distribution of e-values and (**B**) distribution of peptide lengths for the retained epitope mappings.

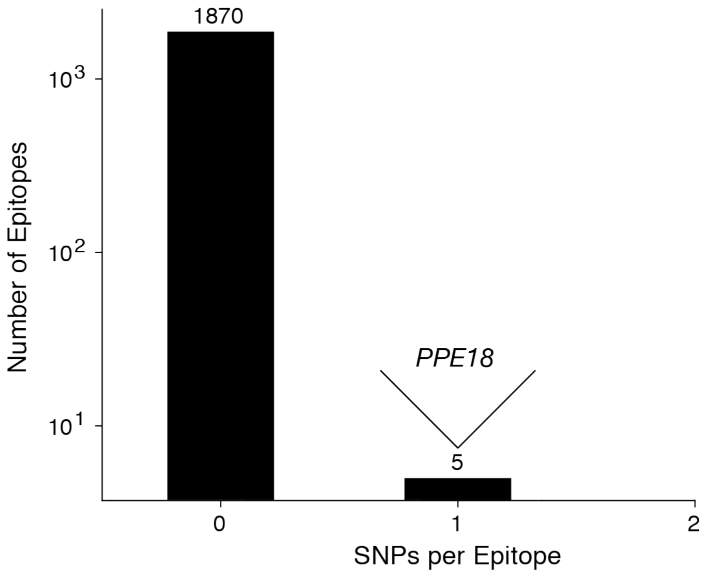


**Fig. S4. Most T cell epitopes remain conserved in-host during active TB disease.** No SNPs were detected in-host for a vast majority of $CD4^+$ and $CD8^+$ T cell epitopes, however 1 SNP was detected in a small number $(n = 5)$ of overlapping epitopes in PPE18. A list of these epitopes is given in **Table S12**.

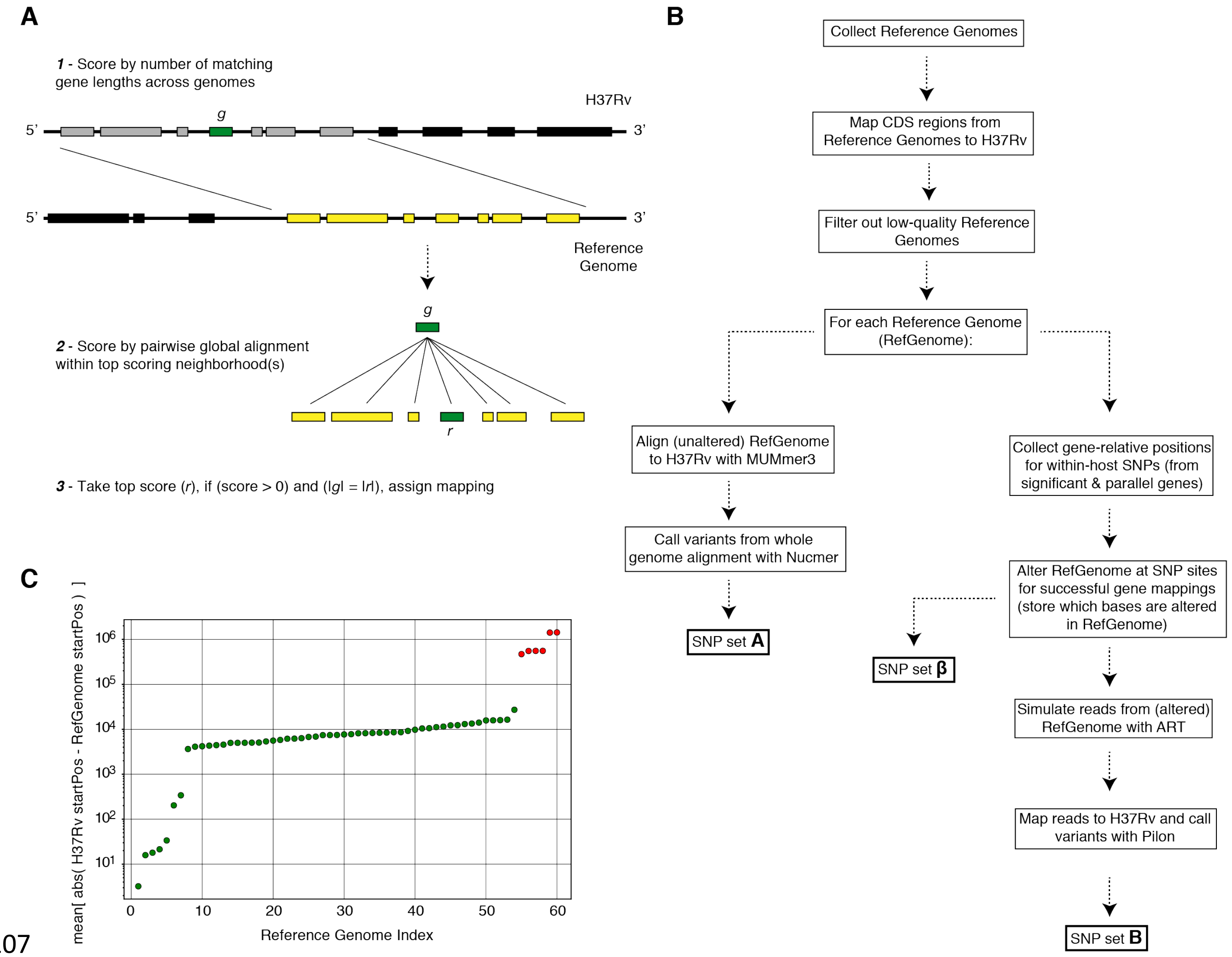


**Fig. S5. Overview of simulation methodology.** To test the accuracy of calling SNPs in repetitive regions with our workflow, we introduced mutations into complete *Mycobacterium tuberculosis* genomes (Reference Genomes), simulated reads from those genomes and assessed the accuracy recalling the mutations from the simulated reads while not introducing spurious mutations (**Supplementary Note 1**). (**A**) We used a sliding window of gene lengths along with a local alignment algorithm to map genes from the H37Rv reference genome to the set Reference Genomes. (**C**) We discarded Reference Genomes that mapped poorly (gene-to-gene) to the H37Rv reference genome (green-RefGenomes kept for simulations, red-discarded RefGenomes).

(**B**) A schematic of our simulation methodology from Reference Genome collection to obtaining SNP sets $\boldsymbol{A}$, $\boldsymbol{B}$ and $\boldsymbol{\beta}$ which are used in our calculations of true positive and false positive calls for each gene (**Supplementary Note 1**).

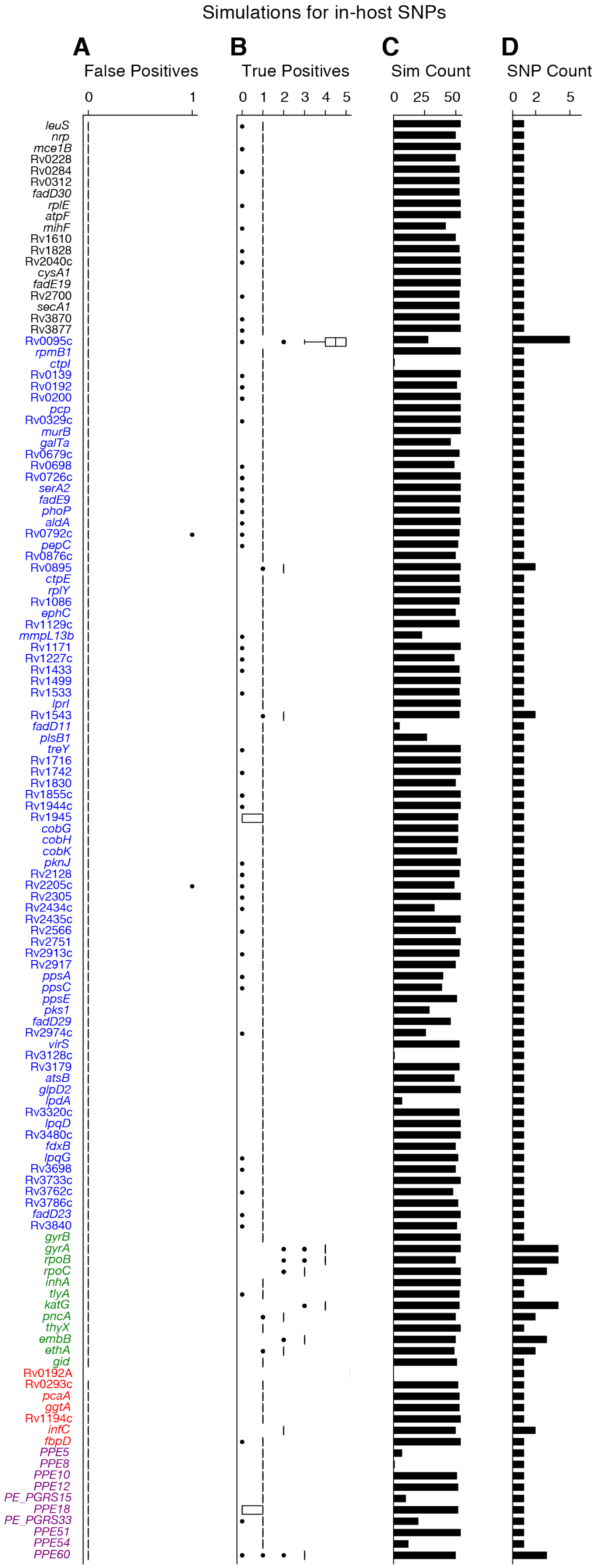
Simulations for in-host SNPs
A
False Positives
0
1
B
True Positives
0 1 2 3 4 5
C
Sim Count
0 25 50
D
SNP Count
0 2 5
leuS
nrp
mce1B
Rv0228
Rv0284
Rv0312
fadD30
rplE
atpF
mihF
Rv1610
Rv1828
Rv2040c
cysA1
fadE19
Rv2700
secA1
Rv3870
Rv3877
Rv0095c
rpmB1
ctpI
Rv0139
Rv0192
Rv0200
pcp
Rv0329c
murB
galTa
Rv0679c
Rv0698
Rv0726c
serA2
fadE9
phoP
aldA
Rv0792c
pepC
Rv0876c
Rv0895
ctpE
rplY
Rv1086
ephC
Rv1129c
mmpL13b
Rv1171
Rv1227c
Rv1433
Rv1499
Rv1533
lprI
Rv1543
fadD11
plsB1
treY
Rv1716
Rv1742
Rv1830
Rv1855c
Rv1944c
Rv1945
cobG
cobH
cobK
pknJ
Rv2128
Rv2205c
Rv2305
Rv2434c
Rv2435c
Rv2566
Rv2751
Rv2913c
Rv2917
ppsA
ppsC
ppsE
pks1
fadD29
Rv2974c
virS
Rv3128c
Rv3179
atsB
glpD2
lpdA
Rv3320c
lpqD
Rv3480c
fdxB
lpqG
Rv3698
Rv3733c
Rv3762c
Rv3786c
fadD23
Rv3840
gyrB
gyrA
rpoB
rpoC
inhA
tlyA
katG
pncA
thyX
embB
ethA
gid
Rv0192A
Rv0293c
pcaA
ggtA
Rv1194c
infC
fbpD
PPE5
PPE8
PPE10
PPE12
PE_PGRS15
PPE18
PE_PGRS33
PPE51
PPE54
PPE60

**Fig. S6. Simulations indicate that we can accurately recall most introduced SNPs while rarely making spurious SNP calls.** We tested the number of true and false positives for each gene with detectable in-host SNPs (**Fig. 5D**). For each gene we collected a set of non-redundant *in-host* SNPs (genomic positions at which these SNPs were called) observed across all subjects (**Table S10**), the number of SNPs collected for each gene is given in (**D**). We then introduced these mutations into 54 complete genomes (RefGenomes) (**Table S9**) and simulated reads after introducing the respective mutations (**Supplementary Note 1**). Only genes that were mapped from H37Rv to a given RefGenome were part of the simulation for that RefGenome. (**C**) The number of successful mappings for each gene (i.e. the number of times each gene was part of a simulation). This is also the number of times true and false positive estimates were calculated for each gene (1 estimate / simulation). (**A**) False positive calls were rarely made across all genes and simulation runs indicating the rarity of false positive SNP calls (calling a mutation that wasn't introduced) made by our pipeline for observed in-host SNPs, even in repetitive regions. (**B**) The number of true positive calls across all genes (across most simulation runs) closely matched the number of introduced SNPs for each gene indicating the rarity of False Negative SNP calls (not calling a mutation that was introduced). We note that no true or false positive estimates for *Rv0192A* were computed since this gene did not map to H37Rv for any of the 54 Reference Genomes used for the simulations.

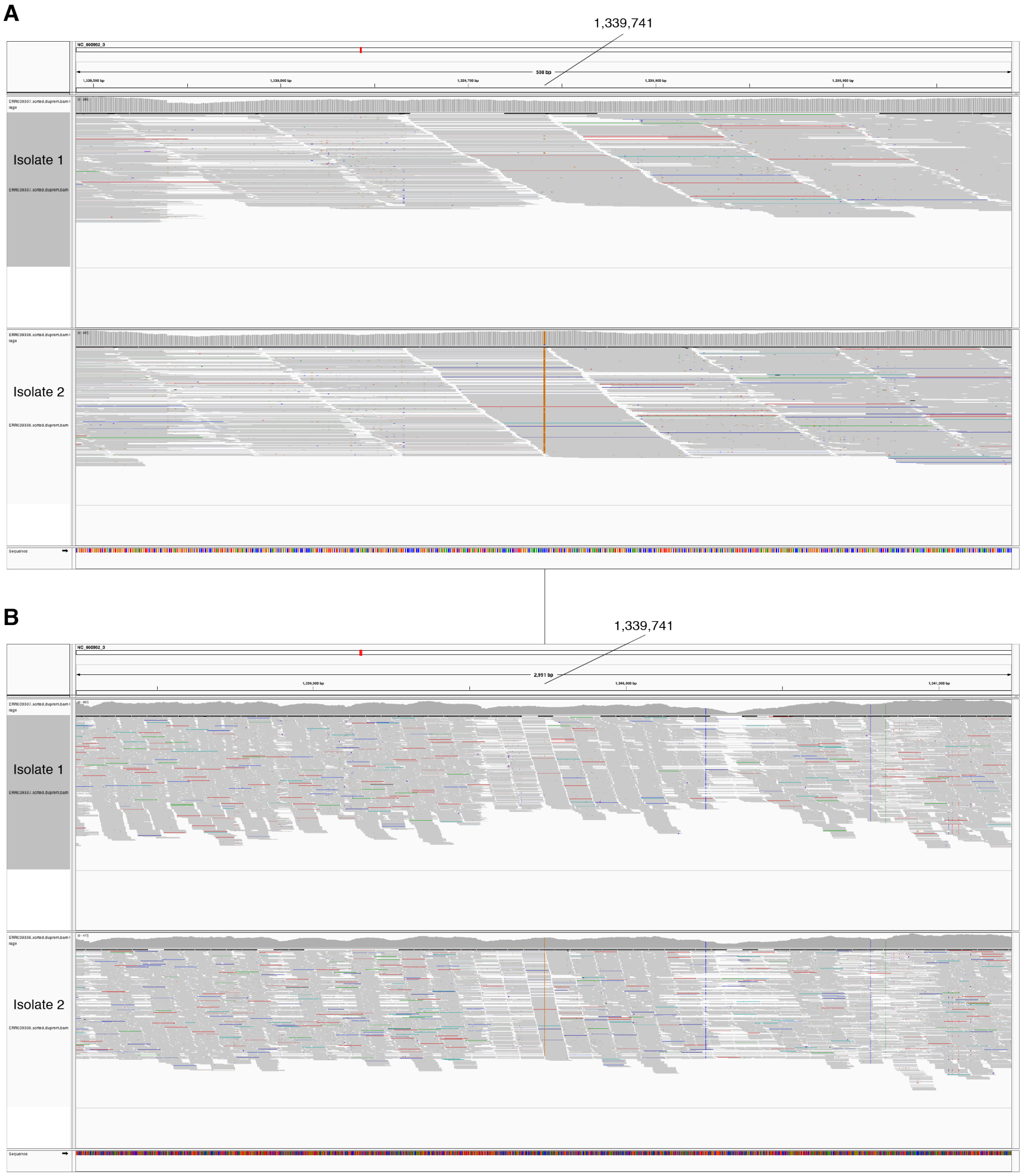


**Fig. S7. In-host SNP detected in *PPE18*.** IGV (Thorvaldsdóttir et al. 2013) image of BAM alignment (reads sorted by start location) for longitudinal clinical isolates that were cultured from sputum collected from patient P000183 (Walker et al. 2013). (**A**) 500 and (**B**) 3000 basepair windows centered at reference position 1339741. Isolate 1 is the BAM alignment for the isolate collected in 2003 and the reference position 1339741 matches the reference allele (C). Isolate 2 is the BAM alignment for isolate collected in 2008 and reference position 1339741 supports an alternate allele (G).

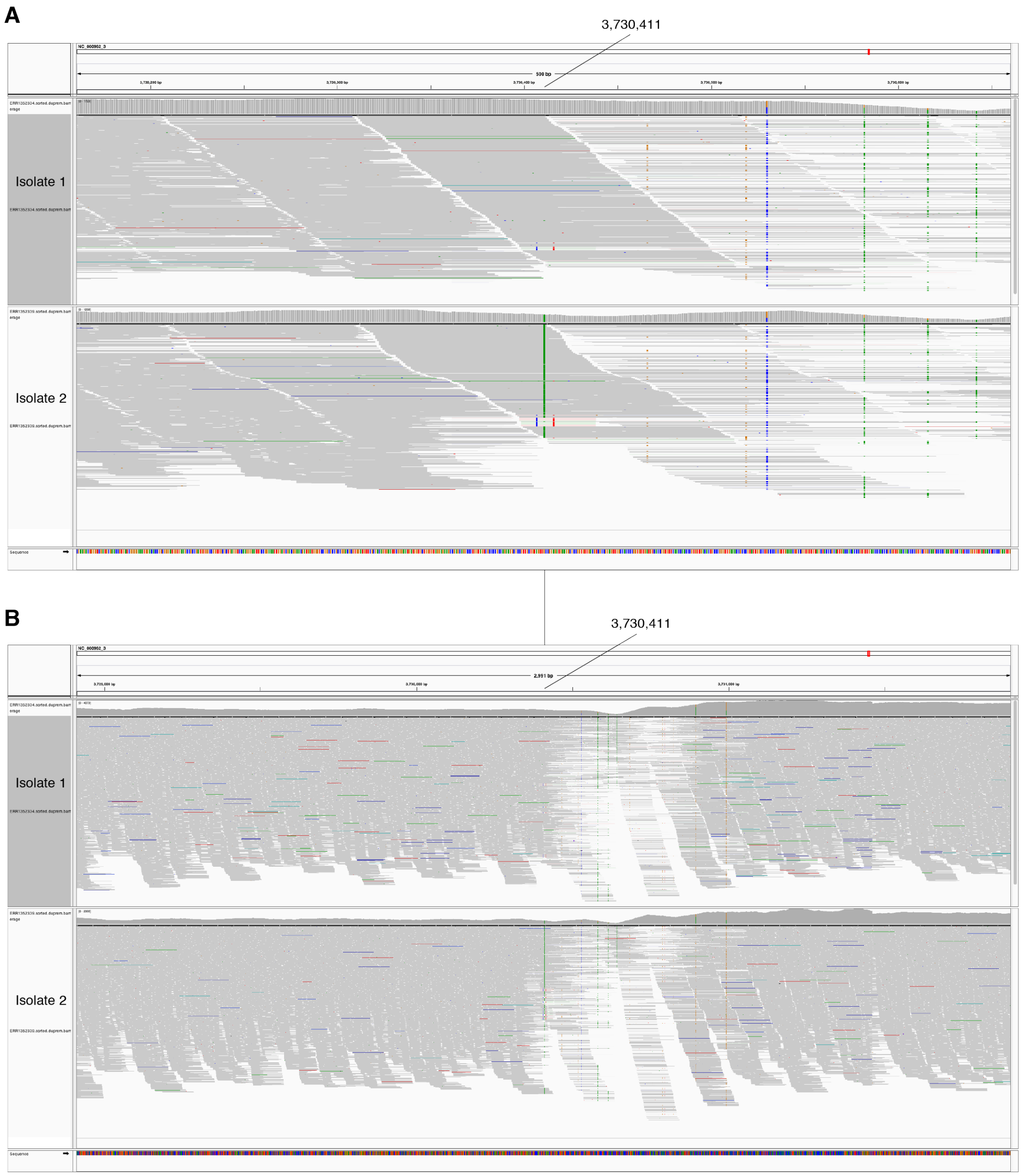


**Fig. S8. In-host SNP detected in *PPE54*.** IGV (Thorvaldsdóttir et al. 2013) image of BAM alignment (reads sorted by start location) for longitudinal clinical isolates that were cultured from sputum collected from patient P09 (Trauner et al. 2017). (**A**) 500 and (**B**) 3000 basepair windows centered at reference position 3730411. Isolate 1 is the BAM alignment for the isolate collected first and the reference position 3730411 matches the reference allele (G). Isolate 2 is the BAM alignment for isolate collected 24 weeks after isolate 1 and reference position 3730411 supports an alternate allele (A).

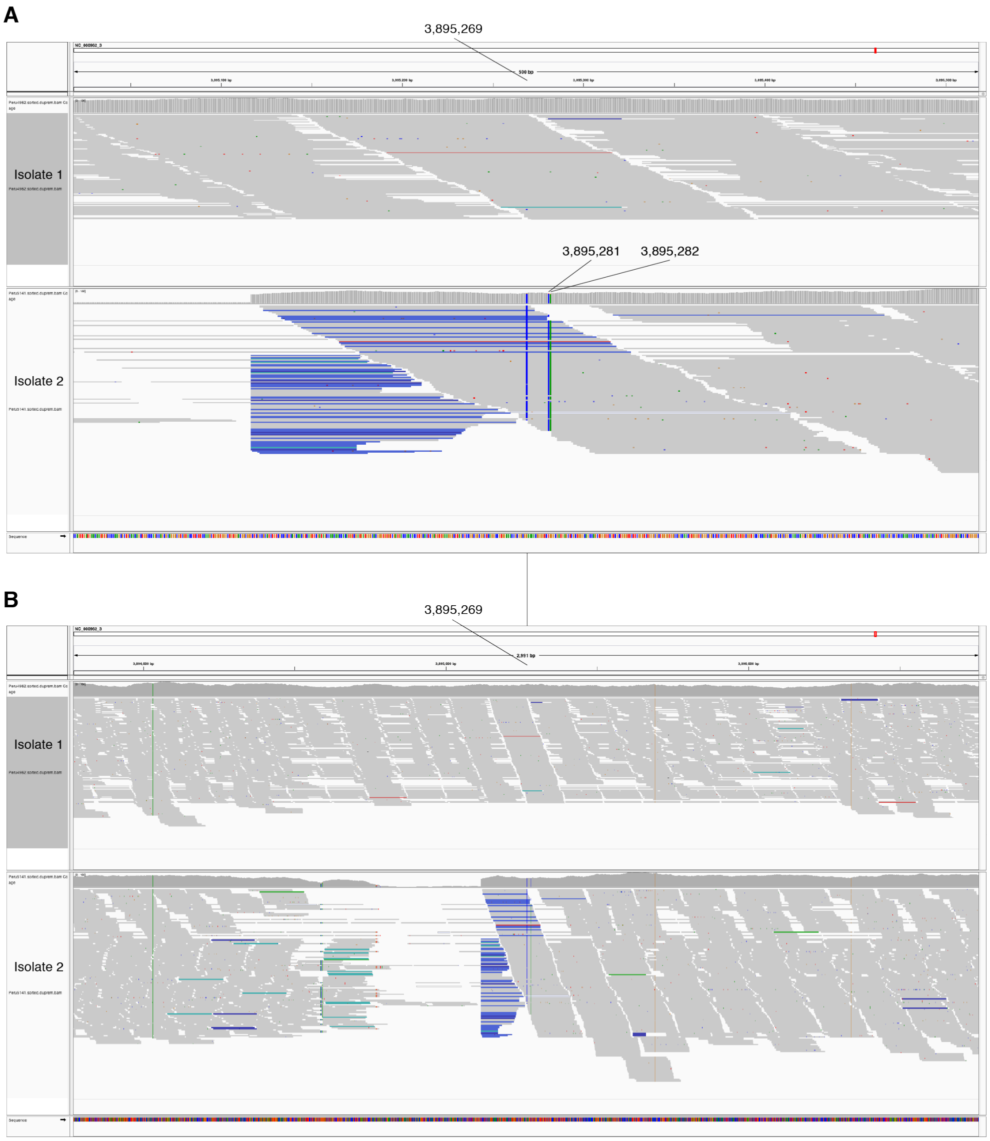


**Fig. S9. In-host SNPs detected in *PPE60*.** IGV (Thorvaldsdóttir et al. 2013) image of BAM alignment (reads sorted by start location) for longitudinal clinical isolates that were cultured from sputum collected from patient 3096 (Farhat et al. 2019). (**A**) 500 and (**B**) 3000 basepair windows centered at reference position 3895269. Isolate 1 is the BAM alignment for the isolate collected on September 25, 2001 and the reference positions 3895269, 3895281, 3895282 match the reference alleles (G,T,G respectively). Isolate 2 is the BAM alignment for isolate collected on October 18, 2002 and reference positions 3895269, 3895281, 3895282 support alternate alleles (C,C,A respectively).

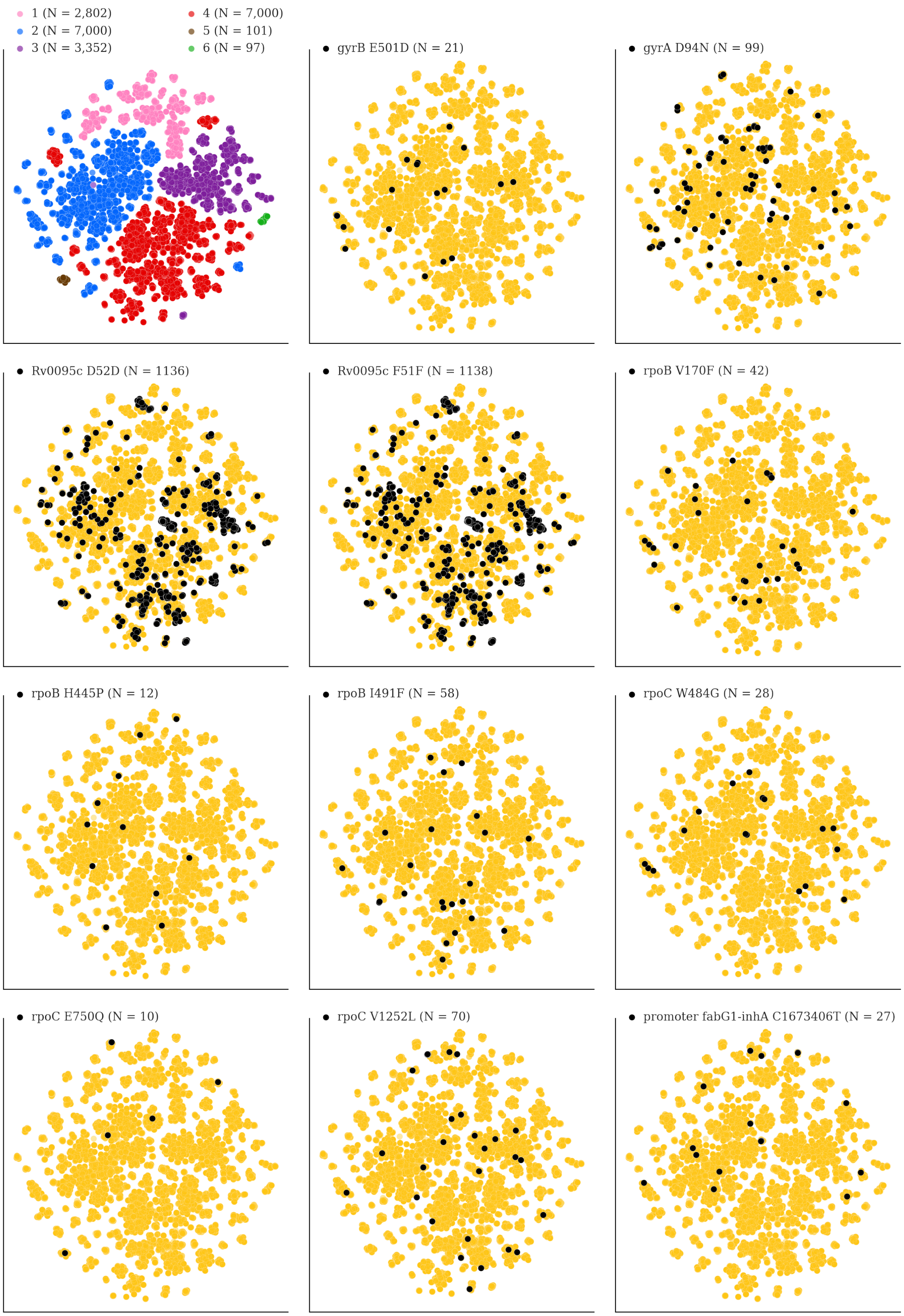


**Fig. S10. Mutations acquired in-host are phylogenetically convergent.** We constructed t-SNE plots from a pairwise SNP distance matrix for our global sample of 20,352 clinical isolates and

128,898 SNP sites (**Methods**). Isolates are colored by global lineage in the first plot, the rest are colored by whether isolates had specific mutations in that were detected in-host (**Table S8**). These mutations occur in a global collection of isolates (**Table S18**) and are scattered across the tSNE plots, indicating that they belong to genetically different clusters of isolates (**Table S19**) and have arisen independently in different genetic backgrounds. Each plot is labeled with the gene name each mutation occurs within, amino acid encoded by the reference allele, H37Rv codon position, and amino acid encoded by the mutant allele (for intergenic mutations - reference allele, H37Rv genome coordinate, and mutant allele). N = number of isolates with mutant allele.

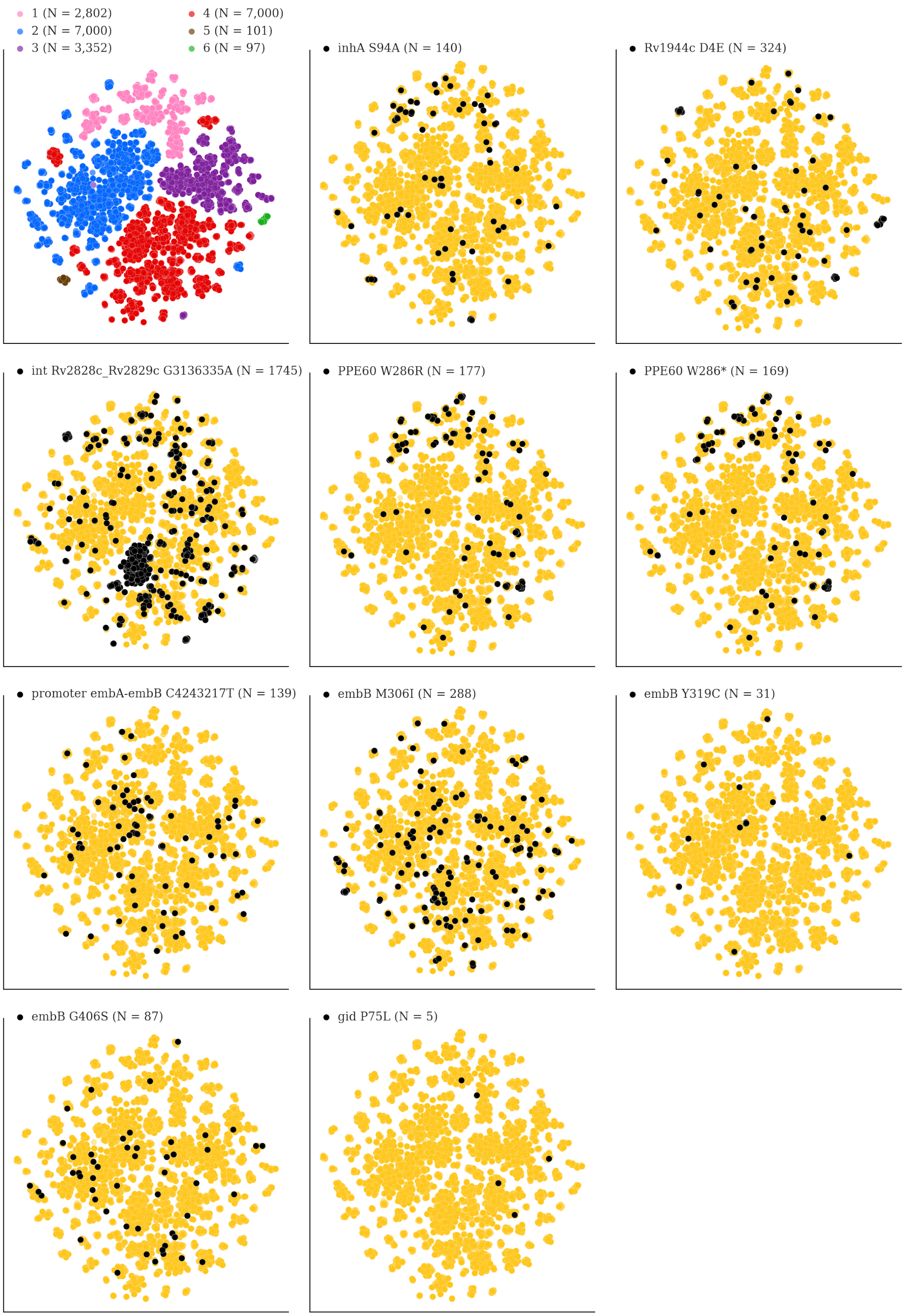


**Fig. S11. Mutations acquired in-host are phylogenetically convergent.** We constructed t-SNE plots from a pairwise SNP distance matrix for our global sample of 20,352 clinical isolates and

128,898 SNP sites (**Methods**). Isolates are colored by global lineage in the first plot, the rest are colored by whether isolates had specific mutations in that were detected in-host (**Table S8**). These mutations occur in a global collection of isolates (**Table S18**) and are scattered across the tSNE plots, indicating that they belong to genetically different clusters of isolates (**Table S19**) and have arisen independently in different genetic backgrounds. Each plot is labeled with the gene name each mutation occurs within, amino acid encoded by the reference allele, H37Rv codon position, and amino acid encoded by the mutant allele (for intergenic mutations - reference allele, H37Rv genome coordinate, and mutant allele). N = number of isolates with mutant allele.

### SUPPLEMENTARY TABLE DESCRIPTIONS

**Table S1.** A table containing details for the eight studies; the sources for the longitudinal isolate pairs. Information includes: (1) reference for each source study, (2) number of subjects included in this study, (3) a description of the sample collection, (4) timing of when sputum samples were collected relative to treatment initiation/cessation (if available).

**Table S2.** A table containing details for all replicate and longitudinal isolates before Kraken, F2, or pairwise SNP filtering.

**Table S3.** A table containing details for all $(n = 400)$ longitudinal isolates used for in-host analysis after filtering for contaminated & mixed isolate pairs.

**Table S4.** A table with the gene categories assigned to each H37Rv locus tag.

**Table S5.** A table containing a list of genomic regions (with H37Rv coordinates) associated with antibiotic resistance.

**Table S6.** A table containing all SNPs (with $\Delta AF \geq 5\%$) in loci associated with antibiotic resistance (**Table S5**) across our sample of 200 longitudinal isolate pairs.

**Table S7.** A table containing all pre-existing antibiotic resistant SNPs detected in the $1^{st}$ isolate collected from each subject with collection dates $\geq 2$ months apart.

**Table S8.** A table containing information for all 174 in-host SNPs detected across all longitudinal isolate pairs.

**Table S9.** A table with details for the 54 publicly available completed (reference) genomes used in our simulations.

**Table S10.** A table with the non-redundant *in-host* SNPs identified within genes and used for SNP calling simulations.

**Table S11.** A table containing all of the epitopes downloaded from IEDB on May 23, 2018.

**Table S12.** A table containing the epitopes belonging to *PPE18* where an in-host SNP was detected.

**Table S13.** A table of all genes identified as *dense*, along with assigned gene category and p-value from mutation density test.

**Table S14.** A table of all genes identified as *convergent*, along with assigned gene category and the number of subjects with an in-host SNP in each gene.

**Table S15.** A table containing the downloaded SEED annotation for H37Rv.

**Table S16.** A table containing the list of H37Rv locus tags corresponding to each subsystem classified by SEED.

**Table S17.** A table containing the pathways and (corresponding in-host SNPs) displaying evidence of parallel evolution.

**Table S18.** A table with details for all SNP calls made in a global collection of 20,352 publicly available isolates after screening for in-host SNPs (**Table S8**).

**Table S19.** A table with details for in-host SNPs (**Table S8**) that displayed a signature of phylogenetic convergence after screening a global collection of 20,352 publicly available isolates (**Methods)**. The number of isolates with each unique mutation (broken down by global lineage) is given.

**Table S20.** A table containing details for isolates that underwent Illumina and PacBio sequencing.
